# Discovery and Functional Characterization of Pro-growth Enhancers in Human Cancer Cells

**DOI:** 10.1101/2021.02.04.429675

**Authors:** Poshen B. Chen, Patrick C. Fiaux, Bin Li, Kai Zhang, Naoki Kubo, Shan Jiang, Rong Hu, Sihan Wu, Mengchi Wang, Wei Wang, Graham McVicker, Paul S. Mischel, Bing Ren

## Abstract

Precision medicine depends critically on developing treatment strategies that can selectively target cancer cells with minimal adverse effects. Identifying unique transcriptional regulators of oncogenic signaling, and targeting cancer-cell-specific enhancers that may be active only in specific tumor cell lineages, could provide the necessary high specificity, but a scarcity of functionally validated enhancers in cancer cells presents a significant hurdle to this strategy. We address this limitation by carrying out large-scale functional screens for pro-growth enhancers using highly multiplexed CRISPR-based perturbation and sequencing in multiple cancer cell lines. We used this strategy to identify 488 pro-growth enhancers in a colorectal cancer cell line and 22 functional enhancers for the *MYC* and *MYB* key oncogenes in an additional nine cancer cell lines. The majority of pro-growth enhancers are accessible and presumably active only in cancer cells but not in normal tissues, and are enriched for elements associated with poor prognosis in colorectal cancer. We further identify master transcriptional regulators and demonstrate that the cancer pro-growth enhancers are modulated by lineage-specific transcription factors acting downstream of growth signaling pathways. Our results uncover context-specific, potentially actionable pro-growth enhancers from cancer cells, yielding insight into altered oncogenic transcription and revealing potential therapeutic targets for cancer treatment.

---

Precise control in gene regulation is the foundation to guide normal cellular functions during various biological processes. Dysregulation of gene expression, especially oncogenes and tumor suppressor genes, contributes to the initiation and maintenance of human cancers (*1–3*). One of the main approaches to cancer therapy is to inhibit the expression of oncogenes selectively. Still, a lack of understanding of these genes’ regulatory mechanisms in cancer cells has been a significant bottleneck. Recent large-scale epigenetic profiling studies, from ENCODE (Encyclopedia of DNA elements), Roadmap Epigenomics Projects, and the Cancer Genome Atlas (TCGA) program, have delineated candidate cis-regulatory elements and cell-type-specific gene regulatory programs in the development and progression of tumorigenesis (*4–10*). The chromatin state and spatial interactions between regulatory sequences and target genes are crucial for cell-type-specific gene expression during normal development (*11–13*). In cancer, abnormal epigenetic changes at regulatory elements are frequently observed and could be used as prognostic biomarkers (*3, 14, 15*). Together, these unique features make enhancers attractive targets in cancer therapy.

One potential solution to achieve selective growth inhibition in cancer cells is to target the cancer-type-specific enhancers that promote cell proliferation (*16–18*). Previous CRISPR-based perturbation screens, using active Cas9 nuclease (Cas9) or nuclease-inactivated dCas9 fused to the Krüppel associated box transcriptional repressor domain (KRAB-dCas9), were performed to identify functional enhancers by deleting or epigenetically silencing candidate genomic regions bearing unique enhancer features, such as DNase-hypersensitive sites (DHS) or H3K27ac (acetylation at the 27th lysine residue of histone H3) or transcription factor binding sites, in one or few cell lines (*19–25*). The limited number of cancer cell types examined in the previous studies did not allow the identification of cancer-type-specific enhancers. Recently, CRISPR-Cas9 based loss-of-function screens were used to identify about 2,000 common essential genes in more than 700 cancer cell lines (*26*). Most of the identified genes were shared across multiple cancer cell types. We hypothesize that cancer-cell-type specific enhancers are involved in the expression of these genes and identification of such pro-growth enhancers could provide ways for selective targeting of specific cancer cell types. To test this hypothesis, we used an unbiased CRISPR interference (CRISPRi) screen to identify pro-growth enhancers in 10 cancer cell lines representing 6 distinct cancer types. We began by focusing on two proto-oncogenes, MYC and MYB. We then carried out a genome-scale CRISPRi screen in one of the cancer cell lines, identified hundreds of pro-growth enhancers and characterized their targets. Furthermore, we found that most pro-growth enhancers are highly accessible in cancer cells but not in primary tissues, and the higher chromatin accessibility of many pro-growth enhancers is associated with poor clinical outcome, suggesting their critical roles in tumorigenesis. We also discovered key upstream factors of the pro-growth enhancers to illuminate gene regulatory program altered in oncogenic transcription.

## Results

### Functional screen of pro-growth enhancers in the *MYC* and *MYB* loci across different cancer cell types

Proto-oncogenes are essential for cell proliferation in normal cells and are frequently activated to promote uncontrollable cell growth during tumorigenesis. The abnormal activation of *c-MYC* proto-oncogene has been implicated in the pathogenesis of most types of human cancer (*27, 28*). Sustained MYC activation is required for tumorigenesis, and the partial suppression of MYC in cancer cells is sufficient to cause acute tumor regression due to the unusual transcriptional addiction in cancer (*2, 29*). MYB, a cell-type-specific oncogene, is important for tumorigenesis in leukemia, colorectal, and breast cancers partially through the regulation of key oncogenes in those cancer types (*30, 31*). Although these oncogenic transcription factors have been well-characterized and are recognized to be critical cancer drivers, it remains challenging to develop small molecules to target them in cancer cells selectively (*32, 33*). As a proof of principle to establish the framework to identify pro-growth enhancers across multiple cancer types, we selected genomic regions near the loci of these two key oncogenes for the screen. We first found that paired guide RNAs (pgRNAs) enable more effective KRAB-dCas9-mediated epigenetic silencing than single guide RNA (sgRNA) (Fig. S1). To generate a universal pgRNA library used in multiple cancer types, we then selected a 3-megabase (Mb) and a 0.6-Mb genomic region that include most known chromatin interactions occurring near *MYC* and *MYB* loci, respectively, in different cell types. Overall, we designed 13,254 tiling pgRNAs targeting those two loci and another set of 1,011 pgRNAs to use as negative controls (non-targeting controls or targeting safe harbor genomic loci). The mean genomic distance between the gRNAs in each pair was ~3 kb, and the genomic spans of adjacent pgRNAs overlapped by 2.7 kb on average (Fig. 1A; Table S1). A previous study (*20*) identified seven functional enhancers (e1-e7) for *MYC* in K562, human chronic myeloid leukemia cells, using CRISPRi with 73,227 sgRNAs targeting all the DHS and genomic regions marked by H3K27ac around *MYC* locus. To examine the performance of our tiling-design strategy, we performed a pooled CRISPRi screen with our pgRNA library in a genetic modified K562 with a fluorescent reporter gene inserted downstream of the *MYC* locus. We carried out the fluorescence-activated cell sorting (FACS) to isolate cells in which MYC expression was reduced and compare the abundance of the pgRNAs in the sorted cells to those in the whole cell population (Fig. S2A-B). To identify the critical enhancers driving *MYC* expression, we used a Generalized Linear Mixed Model (GLMM) framework (*34*) to jointly describe the observed pgRNAs counts across sorted pools under two models: a regulatory model (pgRNA targets a regulatory sequence) and a background model (pgRNA does not target a regulatory sequence).

**Fig. 1.**
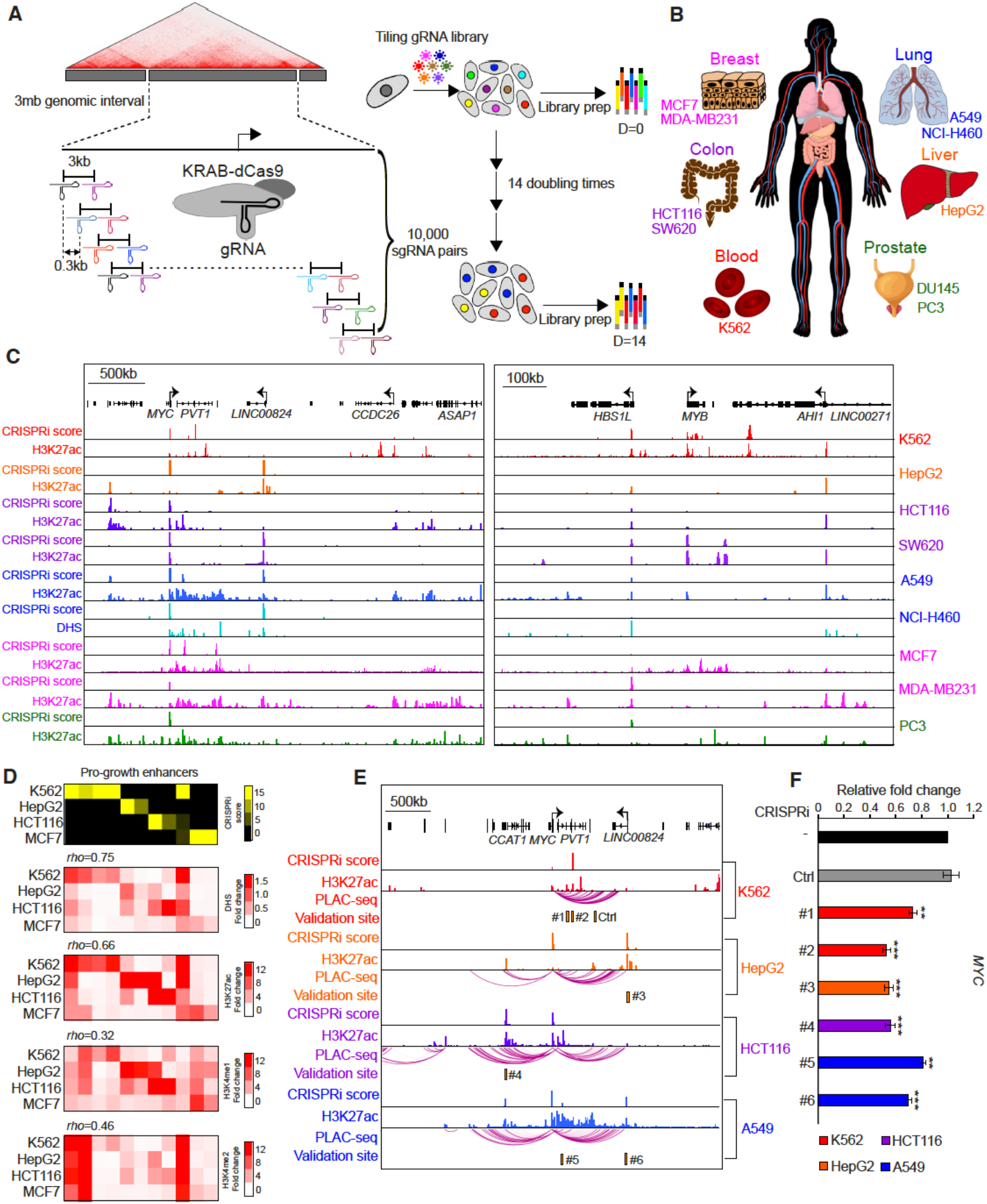
Unbiased CRISPRi screen with a tiling paired sgRNA library to identify pro-growth enhancers around *MYC* and *MYB* oncogenes in major human cancer types. (**A**) Schematic of the experimental strategy illustrates the design of unbiased tiling gRNA screen at the *MYC* locus and the identification of pro-growth enhancers based on cell proliferation assay. (**B**) Major cancer types included in the paired sgRNA CRISPRi screen. CRISPRi screen was carried out in a total of ten different cancer cell lines representing six major cancer types. (**C**) Pro-growth enhancers identified from the screen at the *MYC* and *MYB* loci, with tracks showing CRISPRi score (top) and H3K27ac ChIP-seq signal, H3K27ac (middle), or DNase signal, DHS (bottom), from indicated cell lines. (**D**) Correlation between enhancer features (chromatin accessibility and histone modifications) and the function of pro-growth enhancers across different cell types. (**E**) Chromatin interactions identified by H3K4me3 PLAC-seq at the *MYC* locus in 4 different cancer cell lines. The orange box represents the genomic regions selected for further validation. (**F**) Gene expression measurement of the *MYC* and *MYB* genes by RT-qPCR after silencing the selected pro-growth enhancers by CRISPRi in various cell lines (red: K562; orange: HepG2; purple: HCT116; blue: A549). Data shown are mean ± SD of three technical replicates from one representative experiment of two biological replicates performed. *P*-values were determined by a two-tailed Student’s *t*-test (** and *** indicates *p* < 0.01 and *p* < 0.001 respectively).

As positive controls we used the pgRNAs that target *MYC* promoter to estimate the parameters for the regulatory model and the rest of pgRNAs to estimate parameters for the background model. We then defined a CRISPRi score for a genomic region as the sum of log Bayes factor comparing the two models across pgRNAs overlapping that region (see methods). Using this methodology, we identified five critical *MYC* enhancers with significant CRISPRi scores (> 5) and those enhancers were consistent with the results from previous study (*20*) (Fig. S2C). Our result indicates that this tiling design with pgRNAs has comparable performance to the previously published dense tiling sgRNA design while using 7-fold fewer guide RNA oligos.

Next, we carried out a proliferation-based screen using CRISPRi with the above tiling pgRNA library to identify pro-growth enhancers near the *MYC* and *MYB* loci in 10 different human cancer cell lines representing six major cancer types (lung, breast, liver, colorectal, prostate, and leukemia) (Fig. 1A-B). We compared the abundance of pgRNAs in the initial cell population to the cell population after 14 doubling times and used the same statistical framework to identify candidate pro-growth enhancers (see methods). Together, we identified 11 cancertype-specific and 11 common essential pro-growth enhancers (Fig. 1C and Fig. S3A-B). Further analysis revealed that these pro-growth enhancers were strongly associated with active enhancer marks such as DHS and H3K27ac (Fig. 1D). The majority of these pro-growth enhancers at *MYC* locus were located hundreds of kilobases (kb) away from target oncogenes (Fig. S3C) but exhibited chromatin contacts with the *MYC* promoter, as revealed by Proximity Ligation-Assisted ChIP-seq (PLAC-seq) experiments (*35*) with an H3K4me3 antibody in four cancer cell lines (Fig. 1E). For each of the pro-growth enhancers in the *MYC* locus, we further verified that a significant reduction in *MYC* gene expression is observed after epigenetic silencing pro-growth enhancers by KRAB-dCas9 in different cell types (Fig. 1E-F and Fig. S4).

### Genome-scale identification of pro-growth enhancers in colorectal cancer cells

Aberrant activation of enhancers was previously reported as a key signature in colorectal cancer (*36, 37*). Yet, the function of these enhancers during tumorigenesis is mostly uncharacterized. We selected 532 essential genes, previously identified from a loss-of-function screen in HCT116 colorectal cancer cell line (*38, 39*), and used a pooled CRISPRi screen to interrogate the function of 6,642 distal putative enhancers located near these essential genes together with an additional 4,554 distal candidate enhancers that are associated with H3K27ac mark only in HCT116, but not in the remaining nine cell lines (Fig. 2A). We first selected the top 10 sgRNAs based on the improved criteria and assigned sgRNAs to the same pair based on each sgRNA’s relative genomic location within putative enhancer (see methods). Together, we designed 5 pgRNAs (total 55,980 pgRNAs; Table S2) for each distal putative enhancer to minimize false-negatives due to the variable silencing efficiency of sgRNA (Fig. 2A). Also, we designed 3,520 pgRNAs targeting 460 safe-harbor genomic regions and the promoters of 244 non-expressed genes as negative controls. Using multiple pgRNAs targeting the same distal enhancer facilitated our downstream data analysis using the robust ranking aggregation (RRA) method (*40*) to identify candidate pro-growth enhancers and estimate the false-discovery rate in the screen. Similar design (5-10 sgRNAs targeting TSS or the exon) (*41, 42*) and analysis pipeline (MAGeCK) (*43*) have been developed and widely used to examine gene essentiality in genome-wide CRISPR-Cas9 knockout screens. We first generated several stable KRAB-dCas9 expressed clones of the HCT116 cell line and observed a good correlation in H3K27ac levels with parental cells (Pearson’s *R* = 0.85), indicating that expression of KRAB-dCas9 had minimal effect on the activity of distal enhancers (Fig. S5). We then performed the genome-scale CRISPRi screen in two independent HCT116 clones expressing KRAB-dCas9, and obtained consistent results in the paired gRNA depletion index (Pearson’s *R* = 0.77). As expected, pgRNAs targeting the TSS of essential genes showed a greater reduction in fitness than negative controls pgRNAs (non-targeting or safe-harbor regions) (Fig. 2B). We used the RRA algorithm from MAGeCK (*43*) to obtain the RRA score for individual target genes or putative enhancers. Read counts of negative control pgRNAs were used to estimate the null distribution when calculating the *P*-values. To determine the optimal threshold to select candidate pro-growth enhancers, we estimated the sensitivity and false-positive rate based on the counts of pgRNAs targeting the promoter of 1,085 essential genes as positive controls, and the counts of 3,520 pgRNAs targeting the safe harbor genomic regions and promoters of non-expressed genes as negative controls. Using a threshold corresponding to FDR < 0.2 and 80% sensitivity (Fig. 2C), we identified 558 candidate pro-growth enhancers from 1,338 targets (Fig. 2D; Table S7). We further excluded 70 candidate pro-growth enhancers targeted by multiple pgRNAs with unknown or low specificity scores (see methods; Table S8). We did not observe any enrichment of pgRNAs with low specificity scores among the identified pro-growth enhancers and any significant difference in DNA copy number at pro-growth enhancer loci (Fig. 2E-F), suggesting that the growth defects observed by targeting these enhancers is unlikely due to off-target toxicity.

**Fig. 2.**
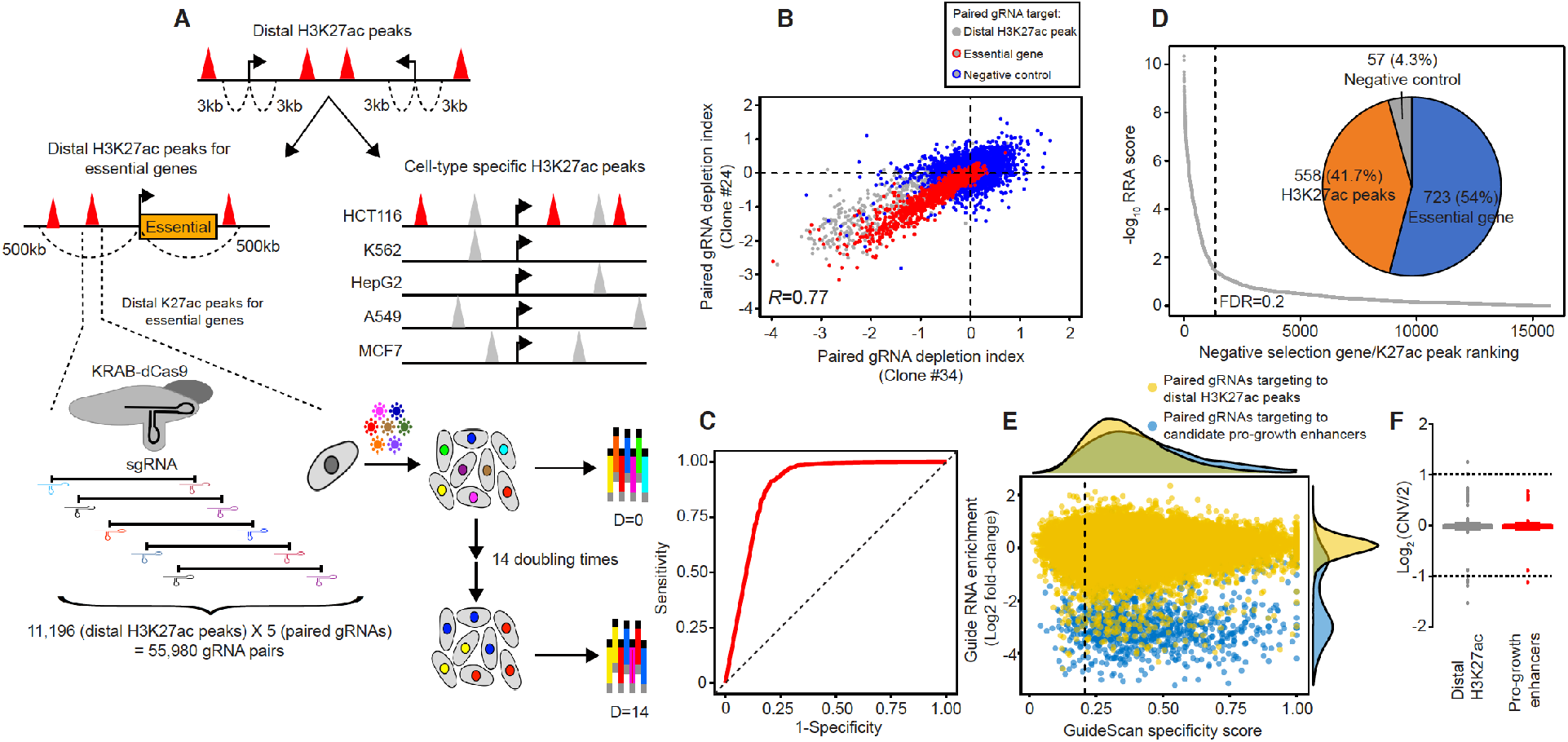
Genome-wide CRISPRi screen to identify pro-growth enhancers in HCT116 colorectal cancer cells. (**A**) Selection criteria for distal putative enhancers to be included in the screen and the design of experimental strategy. The red triangle represents distal H3K27ac peaks selected in this screen, and the gray triangle represents non-cell-type specific H3K27ac peaks. The arrow indicates the transcription start site (TSS). Five paired sgRNAs target each distal H3K27ac peak and the growth effect is measured from a pooled CRISPRi proliferation screen. (**B**) Correlation of fitness effects in two independent KRAB-dCas9 stably expressed clones (Pearson’s *R* = 0.77). The paired gRNA depletion index shown is the average from two biological replicates. Red circled dots indicate paired gRNA targeting positive controls; the TSS of essential genes, and blue circled dots indicate paired gRNA targeting negative controls; safe harbor genomic regions and the TSS of non-expressed gene. Gray dots represent paired gRNA targeting distal H3K27ac peaks. (**C**) The selection of optimal threshold to identify candidate targets from the screen. A receiver operating characteristic (ROC) curve plots the true-positive rate against the false-positive rate for different possible cut-off points in negative selection. (**D**) Pro-growth enhancers identified from the screen. Target with a smaller RRA score (identified by the MAGeCK algorithm) indicates a more substantial negative selection of the corresponding gRNA pairs. A total of 1,338 targets were selected to have significant effects in negative selection (FDR < 0.2; 80% sensitivity). 723 of the significant targets are promoters of essential genes, while 558 are distal H3K27ac peaks, and 57 of them are negative controls. (**E**) Off-target effect assessment using Guidescan score. Comparison of gRNA fitness effect with specificity score. Paired gRNAs targeting pro-growth enhancer are labeled blue, while other paired gRNAs targeting distal H3K27ac peaks are labeled yellow. Low specificity paired gRNAs were considered to those with specificity score below 0.2, indicated by the dashed line. (**F**) Comparison of DNA copy number at all selected distal H3K27ac peaks (N= 11,111) and progrowth enhancers (N= 488). CNV, copy number variation. No significant difference is observed (p = 0.11). *P*-values were determined by two-side Wilcoxon test.

### Pro-growth enhancers regulate cancer cell proliferation in a cell-type-specific fashion

To explore the gene-regulatory program for the 488 pro-growth enhancers identified in HCT116, we predicted their targets by integrating genome-wide maps of chromatin interaction at active gene promoters and scores of gene dependence from previous genome-wide CRISPR-Cas9 knockout screens (Fig. 3A). First, we performed H3K4me3 PLAC-seq assay (*35*) to determine chromatin contacts anchored at active or poised promoters in HCT116 cells. We identified targets of the pro-growth enhancers by if a significant chromatin contact was observed between the promoter and the enhancers using the MAPS software (*44*). In addition, we also searched for possible targets by extending the genomic region 500 kb up or downstream of the pro-growth enhancer that is not paired with target gene using H3K4me3 PLAC-seq data. Subsequently, we intersected the candidate targets with those displaying a significant gene dependence score (>0.7) from the Cancer Dependency Map project (DepMap) (*26*), reasoning that the targets of pro-growth enhancers sought to be also crucial for fitness. Overall, we predicted a total of 910 enhancer and gene pairs (E-G pairs) for 538 pro-growth genes (Table S9). Interestingly, while the vast majority (90.8%) of predicted target genes of pro-growth enhancers were essential genes found commonly across multiple cancer cell lines (*39*) (Fig. 3C), the predicted targets of pro-growth enhancers on average had significantly higher gene expression levels than the rest of common essential genes (Fig. 3B), suggesting pro-growth enhancers further upregulate those predicted pro-growth genes in HCT116 cells. The higher expression of the targets of pro-growth enhancers in HCT116 cells is likely due to cell-type specific enhancers and contacts between the pro-growth enhancers and target promoters, as evidenced by H3K4me3 PLAC-seq data generated in three different cancer types (lung: A549, colorectal: HCT116, leukemia: K562) (Fig. 3D-E). Further supporting this hypothesis, growth defect was observed in HCT116 but not in A549 cells following CRISPRi targeting four of the pro-growth enhancers (Fig. 3F-G), along with the reduction of individual target gene expression selectively in HCT116 (Fig. 3H-I and Fig. S6). These results confirm that the pro-growth enhancers are required for proliferation of cancer cells and the expression of pro-growth gene in a cell-type-specific manner.

**Fig. 3.**
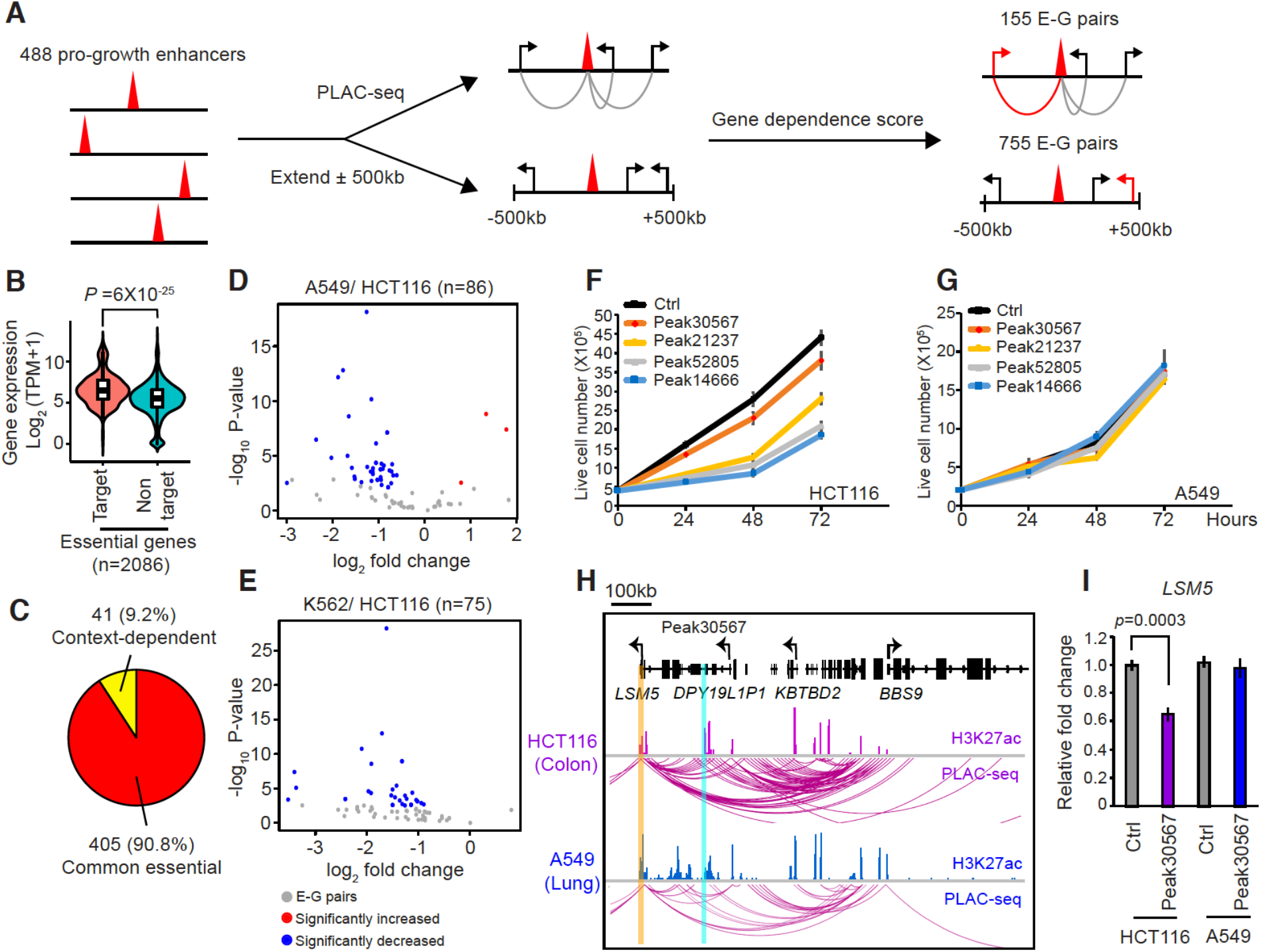
Connecting pro-growth enhancers to target genes with chromatin conformation capture assays. (**A**) Schematic of the *in silico* approach to link pro-growth enhancers to target genes via chromatin interaction and gene dependence. Red triangles represent pro-growth enhancers, and gray arcs indicate chromatin interactions detected by PLAC-seq assays. The arrows indicate the transcription start site (TSS). After the identification of potential target genes, gene dependence is utilized to guide the selection of the predicted target gene (red arrow) and EG pair (red arc). (**B**) Comparison of gene expression in common essential genes identified as targets or non-targets of the pro-growth enhancer. *P* value was determined by the two-sided Wilcoxon test. (**C**) The percentage of context-dependent or common essential genes in the predicted target genes. (**D-E**) Comparison of chromatin contacts between the pro-growth enhancer and target genes in A549 (**D**) with those in K562 (**E**) to HCT116. Significantly increased or decreased chromatin interactions (FDR < 0.05) between the pro-growth enhancer and target gene are labeled in red or blue, respectively. Only E-G pairs with similar H3K4me3 levels at the proximal promoters between A549 and HCT116 (N=86) or K562 and HCT116 (N=75) are used in the differential analysis. (**F-G**) The measurement of cell proliferation after silencing cell-type-specific pro-growth enhancers for HCT116 with KRAB-dCas9 in HCT116 and A549. (**H**) Cell-type-specific chromatin interactions identified by H3K4me3 PLAC-seq in the indicated genomic locus from HCT116 and A549. The pro-growth enhancer and the predicted target gene are highlighted in blue and orange, respectively. (**I**) Relative RNA levels of predicted target gene determined by RT-qPCR after silencing candidate pro-growth enhancer with KRAB-dCas9. Data shown are mean ± SD of three technical replicates from one representative experiment of two biological replicates performed. *P*-value was determined by a two-tailed Student’s *t*-test.

### Pro-growth enhancers are disproportionally associated with poor clinical outcomes

Uncontrolled cell growth through sustained proliferative signaling or evading growth suppressors is the hallmark of cancer (*1*). Our results showed that pro-growth enhancers play critical roles in cell proliferation and regulation of pro-growth genes. To examine whether pro-growth enhancers are selectively active in tumor samples, we analyzed their H3K27ac status in primary tissue from ChIP-seq data generated previously by the Roadmap Epigenomics Project (*45*). We found that most pro-growth enhancers are not associated with H3K27ac in any of the primary tissues examined. As a matter of fact, the number of pro-growth enhancers with H3K27ac ChIP-seq signals in primary tissues is significantly lower compared to that of all the distal candidate enhancers annotated in HCT116 (*P* values < 10^-6^ across 14 tested normal human tissues) (Fig. 4A).

**Fig. 4.**
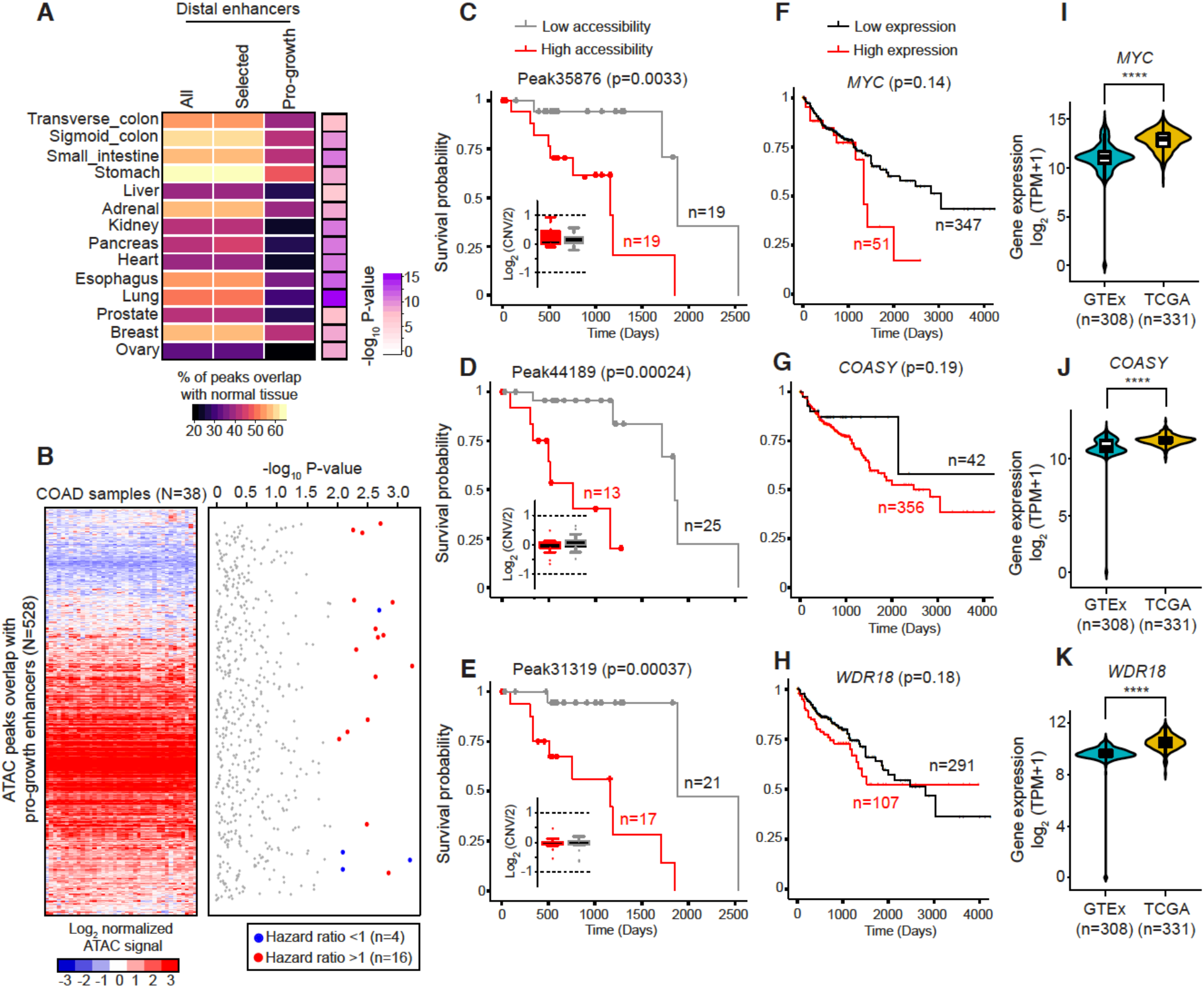
The pro-growth enhancers are enriched for accessible chromatin regions associated with poor clinical outcome of colon cancers. (**A**) Significant depletion of pro-growth enhancers in the genomic regions bearing H3K27ac marks in 14 primary human tissues. Comparison of the percentage of H3K27ac peaks identified from various primary human tissues between all distal H3K27ac peaks (N=22,744), selected distal H3K27ac peaks (N=11,111), and pro-growth enhancers identified in the screen. *P*-values were determined by a hypergeometric test. (**B**) Association of pro-growth enhancer with clinical outcome in colon cancer samples. Left: heat map showing chromatin accessibility of 528 ATAC-seq peaks overlapped with pro-growth enhancers in 38 colon cancer samples. Right: Cox proportional hazards regression analysis of ATAC-seq peaks overlapping with pro-growth enhancers. ATAC-seq peaks with significantly increased or decreased in hazard ratio are labeled as red or blue, respectively (likelihood ratio test; p < 0.01). (**C-H**) Kaplan-Meier analysis of overall survival of colon cancer samples based on averaged chromatin accessibility at ATAC peaks overlapping with pro-growth enhancers (**C-E**) or target gene expression (**F-H**). *P*-values were calculated using the logrank test. In addition, no significant difference is observed in DNA copy number at the indicated pro-growth enhancer loci between high and low accessibility groups (**C-E**). CNV, copy number variation. (**I-K**) Significant increase in gene expression of the pro-growth target at colon cancer samples, obtained from TCGA, compared to normal colon samples, obtained from GTEx (*57*). COAD, colon adenocarcinoma. *P*-values were determined by two-side Wilcoxon test (****: p < 0.0001).

On the basis of our finding that most pro-growth enhancers identified in HCT116 cells are likely cancer cell specific, we hypothesized that their activities in tumor samples might predict clinical outcomes. To test this hypothesis, we examined the chromatin accessibility of the progrowth enhancers in 410 tumor samples across 23 cancer types, mapped previously with the assay for transposase-accessible chromatin using sequencing (ATAC-seq) (*8*). Of the 528 ATAC-seq peaks that overlap with pro-growth enhancers defined in the current study, a significantly higher number (N= 16; *p* = 0.016; one-tailed one sample t-test) were associated with poor clinical outcomes (hazard ratio > 1; *p* < 0.01) than randomly selected ATAC-seq peaks in colorectal cancer samples (Fig. 4B; Fig. S7A). Furthermore, the association between the progrowth enhancer with poor clinical outcome was specific to colorectal cancer. We did not observe a similar increase in hazard ratio, the probability of lower survival rate in the tested group, from HCT116 pro-growth enhancers in breast cancer samples (Fig. S7B), consistent with the role of these pro-growth enhancers in colorectal cancer gene-regulatory programs. Higher chromatin accessibility in pro-growth enhancers correlated significantly with lower survival probability of colorectal cancer patients (Fig. 4C-H). Interestingly, no significant difference in overall survival probability was observed when we used gene expression of the predicted target genes as predictors, despite their elevated expression levels compared to normal counterparts (Fig. 4I-K). These findings indicate that pro-growth enhancers can be utilized as prognostic markers for colorectal cancer.

### Pro-growth enhancers are regulated by lineage-specific transcription factors in cancer cells

Enhancers are regulated by sequence-specific transcription factors (TFs) that interact with them to promote target genes expression in specific cancer cells (*46–48*). To identify the key TFs involved in the regulation of pro-growth enhancers in cancer cell types, we predicted master transcriptional regulators utilizing an integrated computational framework, Taiji (*49*), which estimates the influence of different TFs on gene expression by applying the page-rank algorithm to a model of gene regulatory network based on analysis of transcription factor motifs within the accessible chromatin regions in each cell type or tissue sample. We applied Taiji to previously generated DNase-seq datasets in 262 human cell types/tissues (Table S10) and observed a good correlation (Spearman’s *Rho* > 0.95) of TF ranking scores between similar tissue types (Fig. 5A). Notably, the TFs, which are essential for cell proliferation, also had significantly higher ranking-scores than non-essential TFs (Fig. 5B). Together, these results indicated that key TFs identified by Taiji are master regulators for both gene regulatory program and cell proliferation. To search for the key regulators of pro-growth enhancers, we focused on five cancer cell lines representing five cancer types (leukemia: K562; colorectal: HCT116; liver: HepG2; breast: MCF7; lung: A549) with pro-growth enhancers generated from this study. For the TFs with higher ranking scores in those cancer cell lines, we further selected the TFs with significant context-specific gene dependence (*p* < 0.01) in each cancer type by jointly analyzing the gene knockout data reported by DepMap (*26*) (Fig. 5C; Fig. S8). As expected, this analysis uncovers TCF7L2, a key transcriptional activator of Wnt/ β-catenin signaling pathway (*50*), as a top transcriptional regulator for HCT116 (Fig. 5B; Fig. S8B).

**Fig. 5.**
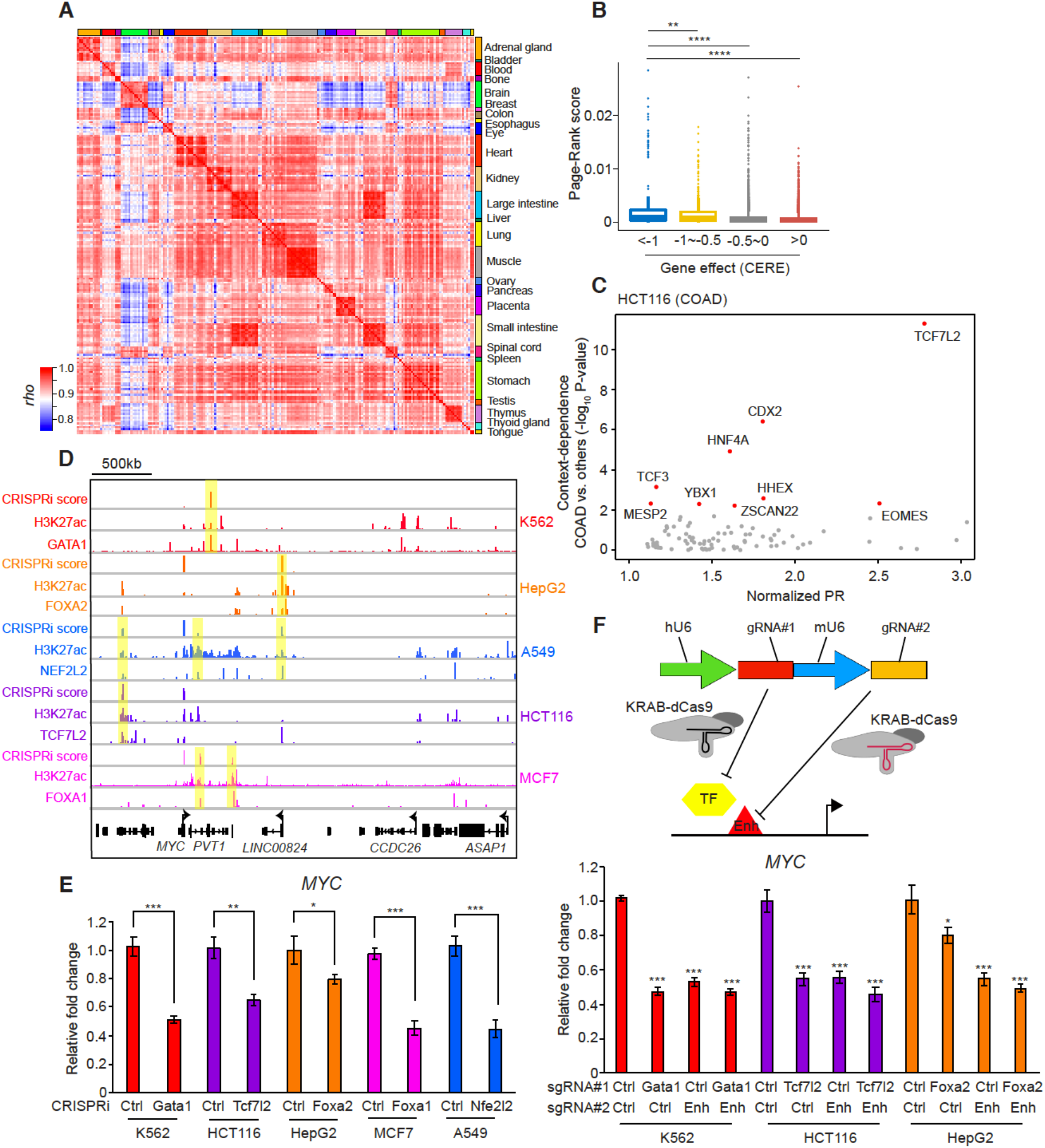
Lineage-specific transcription factors regulate pro-growth enhancers in different cancer cell types. (**A**) Heat map showing pairwise comparison of TF ranking score from Taiji in human primary tissues (n=208). Higher correlation indicates that gene regulatory program is controlled by similar subset of TFs in those tissue types. (**B**) The comparison between TF ranking score and gene dependence (CERE) in 20 DepMap cell lines with available DNase-seq data; each data point represents one TF. *P*-values were determined by two-side Wilcoxon test. (**:*p* < 0.01 and ****:*p* < 0.0001). (**C**) The criteria to select candidate TFs in HCT116 based on TF ranking score and context-specific gene dependence. The red dots represent candidate TFs with high normalized page-rank score and significant context-dependence (p < 0.01). COAD, colon adenocarcinoma. *P*-values were determined by two-side Wilcoxon test. (**D**) ChIP-seq signal of lineage-specific TFs around *MYC* locus overlapping with pro-growth enhancers. Genome browser snapshots show the selective enrichment of GATA1, FOXA2, NEF2L2, TCF7L2, and FOXA1 bindings at pro-growth enhancers identified from K562, HepG2, A549, HCT116, and MCF7, respectively. Pro-growth enhancers with strong enrichment of lineagespecific TF binding are highlighted in yellow. (**E**) Gene expression of *MYC* measured by RT-qPCR after silencing the selected lineage-specific transcription factor with KRAB-dCas9 in various cancer cell lines. Data shown are mean ± SD of three technical replicates from one representative experiment of two biological replicates performed. *P*-value was determined by a two-tailed Student’s *t*-test. (*: *p* < 0.1, **: *p* < 0.01 and ***:*p* < 0.001). (**F**) RT-qPCR for relative RNA levels of *MYC* expression in cells expressing sgRNA pairs targeting individual progrowth enhancer and lineage-specific TF or both. Data shown are mean ± SD of three technical replicates from one representative experiment of two biological replicates performed. *P*-value was determined by a two-tailed Student’s *t*-test. (*:*p* < 0.1 and ***: *p* < 0.001).

To test if TCF7L2 is responsible for pro-growth enhancers in HCT116, we analyzed previous ChIP-seq and RNA-seq experiments (*51, 52*). Gene expression for ninety-five of the predicted targets was significantly downregulated after *Tcf7l2* knockout (Fig. S9A), significantly higher than expected in randomly selected pro-growth enhancers (N= 25) (Fig. S9B). In addition, TCF7L2 bound to 125 (26%) of the pro-growth enhancers in HCT116, selective higher than in other cancer cell types (breast cancer MCF7: 9%; liver cancer HepG2: 7%; pancreatic cancer Panc1: 13%; embryonic kidney HEK293: 6%) (*51*). Furthermore, we showed that lineage-specific TFs were required for the activities of pro-growth enhancers of *MYC* (*53*) in different cancer cell lines including HCT116 (Fig. 5D), as their knock down using CRISPRi resulted in a cell-type-specific reduction of *MYC* gene expression (Fig. 5E). Moreover, CRISPRi of either lineage-specific TFs or pro-growth enhancers alone achieved the same levels of effect on *MYC* expression as silencing both simultaneously, supporting the hypothesis that these TFs worked through pro-growth enhancers to mediate cell-type specific expression of MYC genes (Fig. 5F). These results, taken together, suggested that the lineage specific TFs play a critical regulatory role in activation of pro-growth enhancers and targeting of these factors could achieve selective growth inhibition in a cancer type specific manner.

## Discussion

Despite the rapid progress in identifying putative enhancers in the human genome from recent years, the function of most annotated human enhancers is still untested. Here, we utilized a robust, unbiased, and high throughput functional screen to systematically identify pro-growth enhancers in human cancers. From a genome-scale survey of putative distal enhancers, we identified 488 pro-growth enhancers essential for cancer proliferation, the majority of which seem to be inactive in normal tissues. Additionally, we uncovered a cell-type-specific gene regulatory program for common essential genes, which are generally not considered as therapeutic targets. Our findings suggest that targeting cancer-type-specific enhancers could achieve selective proliferation inhibition for cancer treatment. In addition to their critical roles in promoting proliferation, we also showed that higher chromatin accessibility at pro-growth enhancers is selectively associated with low survival probability and might lead to a malignant state in colorectal cancer. Overall, this systematic functional perturbation assay to identify progrowth enhancers is highly generalizable. It can be readily applied to other cancer cell lines, potentially leading to the discovery of new therapeutic targets in cancer research. Similarly, this strategy could also be adapted to discover enhancers crucial for different physiological phenotypes, such as cellular differentiation, cellular responses to extracellular signaling, etc.

Overexpression of oncogene is a hallmark of many cancer types, but many oncogenes are considered “undruggable” due to their essential role in normal cell growth. Our study identified cell-type-specific pro-growth enhancers, and established a role for lineage-specific TFs as key transcription factors that regulate the pro-growth enhancers in cancer cells. Our finding provides a rationale to target lineage-specific TFs to selectively inhibit cancer cell growth. While it is still difficult to develop selective inhibitors for lineage-specific TFs, we anticipate that several alternative approaches, including peptide nucleic acid, small interfering RNA, or CRISPR system, which have been utilized to target gene promoter (*54*), distal enhancer (*55*), or enhancer RNAs (*56*) and lead to activate or repress target gene expression, could also be applied to target pro-growth enhancers. Together, our study provides the first large-scale map of pro-growth enhancers, reveals cell-type-specific gene regulatory program for common essential genes, and characterizes key upstream regulators of pro-growth in multiple cancer types.

## Acknowledgments

We thank H. Zhao for MYC reporter knock-in constructs; J.S. Weissman for lenti-KRAB-dCas9-BFP plasmid; C.A. Gersbach for lenti-KRAB-dCas9-sgRNA vector; G. Li, Q. Zhu, M. Tastemel, and all members of the Ren lab for many productive and exciting discussions of this work.

## Funding

This study was supported by grant 1UM1HG009402 from the National Human Genome Research Institute (NHGRI);

## Author contributions

P.B.C., S.W., P.S.M. and B.R. designed the study. P.B.C. and R.H. carried out the experiments. P.B.C and B.L. developed computational method to design paired sgRNAs for genome-wide screen. P.B.C, B.L., P.C.F., K.Z., N.K., S.J., G.M. and B.R. contributed to data analysis and interpretation. P.B.C, P.C.F., K.Z, N.K., S.J., B.R. wrote the manuscript with input from all authors. B.R. supervised the work and obtained funding;

## Competing interests

B.R. is a co-founder and consultant of Arima Genomics, Inc., and co-founder of Epigenome Technologies Inc; and P.S.M. is a cofounder and consultant of Boundless Bio, Inc.;

## Data and materials availability

All data is available in the manuscript or the supplementary materials. Sequencing data have been deposited in the Gene Expression Omnibus under accession number GSE161874 and the following are available for use online: RELICS v1 (https://github.com/patfiaux/RELICS/releases/tag/v1.0), CRISPY (https://github.com/MichaelMW/crispy), and MAPS (https://github.com/ijuric/MAPS).

## Supplementary Materials

Materials and Methods

Figures S1–S9

Tables S1-S11

References (*1–81*)

## Materials and Methods

### Cell line

Human chronic myeloid leukemia cell line K562 was cultured in RPMI with 10% fetal bovine serum (FBS) (Gemini). Human liver cancer cell line HepG2, colon cancer cell line SW620, lung cancer cell line A549 and NCI-H460, breast cancer cell line MDA-MB231, prostate cancer cell line DU-145 and PC3 were cultured in DMEM/F12 media (Thermo Fisher) with 10% FBS. Human colorectal cancer cell line HCT116 was cultured in McCoy’s 5A Modified media (Thermo Fisher) with 10% FBS, and human breast cancer cell line MCF7 was cultured in DMEM (4500mg/L glucose) (Thermo Fisher) with 10% FBS. All human cancer cell lines were obtained from ATCC and tested negative for mycoplasma.

### Antibodies

Antibodies used in this study were mouse anti-H3K27ac (Diagenode, C15200184) and rabbit anti-H3K4me3 (Millipore, 04-745).

### Chemical inhibitors

Chemical inhibitors used in this study were SNS-032 (Selleckchem, S1145) and Apicidin (Sigma, A8851).

### MYC reporter knock-in line generation

K562 cells were electroporated with CRISPR expression plasmids, and donor constructs from the previous study (*58*) using cell line nucleofector Kit V (Lonza) following the manufacturer’s instructions. Cells were selected by puromycin (2 μg/ml) (InvivoGen) 3 days postelectroporation. After seven day selection, GFP positive cells were identified and isolated using a SH800S cell sorter (Sony) to one cell per well in a 96-well plate. The purity of the individual clone was examined by FACS and genotyped by PCR.

### CRISPRi line generation

HCT116 cells were transduced with lentivirus carrying KRAB-dCas9-BFP (Addgene #85969). Three days after transduction, BFP positive cells were isolated using a SH800S cell sorter (Sony) to one cell per well in a 96-well plate. The purity of the individual clones was examined by FACS, and several clones with the strongest BFP signal were selected for the following experiment.

### Selection of genomic targets for tiling CRISPRi screen

In the pilot study, we selected two key oncogenes, *MYC* and *MYB*, that are required for cellular proliferation in various cancer types. We followed a similar principle as the previous study (*20*) to identify genomic regions for the tiling CRISPRi screen. Briefly, chromatin organization is known to play key roles in gene regulation, and the majority of enhancer and promoter interactions occur within topological associated domains (TAD) (*12, 59*). Because cell-type-specific chromatin interactions occur around the *MYC* locus, we combined Hi-C data from K562, HepG2, A549, HCT116, and selected genomic regions to include the entire TAD around *MYC* (3-Mb) and *MYB* (~600 kb) loci.

### Selection of genomic targets for genome-wide CRISPRi screen

We first identified pro-growth enhancers sharing similar epigenetic features as active enhancers from the pilot screen. That finding provided us the rationale to focus on H3K27ac peaks instead of an unbiased screen, which is more labor and cost-intensive on a genome-wide scale setting. Second, our data supported that chromatin interactions between pro-growth enhancers and their targets are important for gene regulation. Thus, we further narrowed down to those H3K27ac peaks located within 500-kbp around target genes because promoter capture Hi-C data have shown that most three-dimensional (3D) promoter-based interactions occur within a 500-kbp distance (*60, 61*). Third, we were able to validate several pro-growth enhancers identified from the pilot screen, indicating cell fitness is a reliable readout to reflect the reduction in gene expression of essential genes. However, we are conscious of the possible limitation in this screen’s sensitivity to identify weak pro-growth enhancers, which only slightly or moderate regulate target gene expression, from the proliferation-based screen. Last, our screen identified several context-dependent pro-growth enhancers, indicating that cell-type-specific H3K27ac peak is the potential candidate of pro-growth enhancer. Overall, we selected 6,642 H3K27ac peaks based on genomic location and 4,554 H3K27ac peaks based on cell-type-specific activity.

### sgRNA design for tiling sgRNA library

We used the guide RNA design tool, developed in the previous study (*62*), to identify unique sgRNAs and select sgRNA pairs in two genomic loci (*MYB* locus — chr6: 134,923,863-135,478,863; *MYC* locus — chr8: 127,182,756-130,337,754). For the selected genomic loci, each sgRNA pair’s averaged distance was 3-kbp with 0.3-kbp step size. We generated 11,681 and 1,573 sgRNA pairs for *MYC* and *MYB* loci, respectively. We also included 955 sgRNA pairs that lacked the PAM sequence targeting MYC locus and 56 sgRNA pairs targeting safe harbor regions as negative controls and 170 positive sgRNA pairs targeting essential genes (*Gata1* and *Phb*) and previously identified *MYC* enhancer in K562 cells (e1-e7). sgRNA information is listed in Table S1.

### sgRNA design for genome-wide sgRNA library

We included 11,196 distal H3K27ac peaks and designed five sgRNA pairs targeting each peak. To maximize the coverage of the sgRNA pair, we assigned sgRNAs to the same pair based on the relative location within each H3K27ac peak. For instance, we paired the first sgRNA closest to the start of the H3K27ac peak with the sixth sgRNA closest to the start of the same H3K27ac peak. In general, we selected the top 10 sgRNAs for each H3K27ac peak based on the improved criteria that we learned from the pilot tiling screen. Briefly, we found that ~40-60% sgRNA pairs located within validated functional enhancers are effective with higher on-target score (*63*). Furthermore, we observed the percentage of effective sgRNAs increasing to ~60-90% when we only considered sgRNAs identified from our developed sgRNA design (*62*) and FlashFry tools (*64*). To minimize potential false-negatives due to low sgRNA efficiency, we selected the top 10 sgRNAs, based on the improved selection criteria, targeting for each distal H3K27ac peak included in the screen. In addition, we used the same approach to design 2,300 sgRNA pairs targeting 460 safe harbor genomic regions as negative controls. To assess genome-wide screen performance, we also designed five paired sgRNAs targeting 1,018 essential genes using sgRNA rank score from previous CRISPRi genome-wide screen study (*65*). Overall, we generated a genome-wide sgRNA library containing 55,980 sgRNA pairs targeting 11,196 distal H3K27ac peaks, 5,391 sgRNA pairs targeting essential genes as positive controls, and 3,520 sgRNA pairs targeting safe harbor, promoter of non- or low-expressed gene, and non-targeting control as negative controls. sgRNA information is listed in Table S2.

### Tiling and genome-wide CRISPRi plasmid library construction

We designed, synthesized the pool of paired-guide RNA oligo (Agilent), and generated the dual-gRNA plasmid library as previously described (*62*) with the following modifications. Briefly, we amplified the oligo library by PCR with less than 20 cycles. After PCR, the pooled oligo library was purified with AMPure XP beads (Beckman Coulter). The lentiviral vector carrying dCas9-KRAB was obtained from Addgene (Plasmid #71236) and linearized with BsmBI followed by gel purification. We performed 5 or 20 Gibson assembly reactions for tiling or genome-wide sgRNA library followed by manufacturer’s instructions (NEB) and purified DNA using ethanol precipitation. The purified DNA was electroporated into Endura competent cells (Lucigen) using 50-100ng DNA per electroporation (we set up 6 or 24 replicate transformation for tiling or genome-wide sgRNA library to maintain library complexity), and the colonies were harvested within 14 hours at 30°C to minimize recombination activity in bacteria. We extracted the plasmids using the Plasmid Maxi prep kit (Macherey-Nagel).

### Lentivirus generation

Briefly, 5ug of plasmid library was co-transfected with four ug PsPAX2 (Addgene #12260) and one ug pMD2.G (Addgene #12259) into a 10-cm dish of 293FT cells (Thermo Fisher) in DMEM containing 10% FBS using FuGene HD (Promega). Scale up the number of 293FT cells and transfection depending on the yield and library size. The growth medium was replaced 12 hours after transfection, and the lentivirus was harvested 48 hours post-transfection. The viral titer was determined for each individual cancer cell line using the survival cell number under puromycin selection from a serial dilution of lentivirus transduction.

### Pooled CRISPRi screens for essentiality

We infected cells with lentiviral libraries at a low multiplicity of infection (MOI=0.5) to ensure each infected cell got one viral particle. In general, we maintained cell numbers with at least 1000 fold coverage of the lentiviral library during the entire proliferation screen. For instance, we would have at least 15 million survived cells after puromycin selection for a pooled library with 15,000 paired-guide RNAs. We started puromycin selection (2 μg/ml) (InvivoGen) 48 hours post-transduction for at least two days or until no survival cell was observed from the control group. After puromycin selection, we recovered and cultured cells in a lower puromycin concentration (0.2 μg/ml) for additional two days. To start the screen, we collected at least 15 million cells as “doubling time 0” and sub-cultured at least 15 million cells for 14 doubling times in a lower concentration of puromycin (0.2 μg/ml). The cell concentration, viability, and doubling time were examined every two days. We sub-cultured and split cells when they reached more than 90 % confluency. In the end, at least 15 million cells that reached 14 doubling times were harvested. For the proliferation screen, we performed two replicate experiments for every human cancer cell line.

### Sorting-based CRISPR and CRISPRi screen

K562 MYC reporter cells were transduced with lentivirus carrying Cas9 or KRAB-dCas9 with tiling sgRNA pairs pooled library using MOI=0.3. Forty-eight hours post-transduction, cells were selected using blasticidin (8 μg/ml) (Thermo Fisher) for nine days. The cells were split in a 1 to 4 ratio every three days. We used at least 60 million survived cells after blasticidin selection for a pooled library with 15,000 paired-guide RNAs to start the screen. Cells were sorted into six different bins (Bin1-6) based on GFP signals using a SH800S cell sorter (Sony).

### Generation of Illumina sequencing library

Genomic DNA was isolated from proliferation-based (D=0 and D=14) or sorting-based (Bin #1-#6) screens and used to generate illumina sequencing libraries. Briefly, we used 400 ng genomic DNA as a template per PCR reaction. In 1^st^ PCR, the library was PCR amplified using Herculase II (Agilent) in the 96-well plate to increase the coverage for 22 cycles with the following program (98°C for 5 min; 98°C for 35 sec, 52°C for 30 sec, 72°C for 1min and repeat for 21 cycles; 72°C for 5 min). After 1^st^ PCR, we combined all reactions from the 96-well plate and used 2 ul of the mixture as the 2^nd^ PCR amplification template. In 2^nd^ PCR amplification, we amplified the library with Truseq index primers and prepared two PCR reactions per library. The 2^nd^ PCR was amplified using KAPA Hi-Fi (KAPA bioscience) for five cycles with the following program (95°C for 3 min; 98°C for 20 sec, 65°C for 15 sec, 72°C for 30 sec and repeat for four cycles; 72°C for 1 min). The sequencing library was combined and gel-purified (~690bp). We generated 100 bp paired-end reads on Illumina Hiseq 4000. Primer information for illumina sequencing library is listed in table S3.

### Cloning individual sgRNAs

The lentiviral vector carrying dCas9-KRAB was obtained from Addgene (Plasmid #71236) and linearized with BsmBI followed by gel purification. sgRNA oligo (Table S4) was annealed and phosphorylated before the ligation. Individual sgRNA construct was verified using Sanger sequencing (Genewiz).

### Cloning paired sgRNAs

sgRNA pair cassette was PCR amplified from the gBlock template (*62*) containing tracRNA and mouse U6 promoter sequence using KAPA Hi-Fi (KAPA bioscience) with primers to add homology arms (Table S5) for Gibson assembly. We assembled a 20 ng amplified cassette into a 50 ng digested vector in a 20 μL Gibson reaction (NEB). The individual construct was verified using Sanger sequencing (Genewiz).

### RNA extraction and quantitative RT-PCR

We used quantitative RT-PCR (RT-qPCR) to validate the effect of selected enhancers in gene regulation. In CRISPRi experiments, cells transduced with lentivirus carrying KRAB-dCas9 (Addgene #71236) and sgRNAs were harvested one week after the transduction. In addition, cells treated with 1 μM of SNS-032 (Selleckchem) or Apicidin (Sigma) were harvested 24 hours after inhibitor treatment. Total RNA was extracted using Trizol (Thermo Fisher) following the manufacturer’s instructions. Reverse transcription was performed for one hour using random priming (Promega). qPCR reactions (0.5 μl cDNA, 0.2 μM each primer, SYBR green Master Mix (Kapa biosystems) were performed on a Roche LightCycler 480 Real-Time PCR detection system, using primers specific for each gene (Table S6). Data were normalized to loading controls (Gapdh).

### Chromatin immunoprecipitation-sequencing (ChIP-seq)

ChIP-seq experiments for H3K27ac mark were performed as described in ENCODE experiment protocols (“Ren Lab ENCODE Chromatin Immunoprecipitation Protocol” in https://www.encodeproject.org/documents/) with minor modificaitons. The cells from ~80% confluent 10 cm dishes were crosslinked by adding fixation solution (1% formaldehyde, 0.1M NaCl, 1 mM EDTA, 50 mM HEPES·KOH pH 7.6) for 10 min at room temperature. Crosslinking was quenched with 125 mM Glycine for 5 min. We used 1.0 million cells for each ChIP sample. Shearing of chromatin was performed using truChIP Chromatin Shearing Reagent Kit (Covaris) according to the manufacturer’s instructions. Covaris M220 was used for sonication with following parameters: 410 seconds duration at 20.0% duty factor, 75.0 peak power, 200 cycles per burst at 5-9°C temperature range. For immunoprecipitation, we used 50 μL Protein A or Protein G Magnetic beads (NEB) and washed twice with PBS with 5 mg/ml BSA and 4 μg of antibody coupled in 500 μl PBS with 5 mg/ml BSA overnight at 4°C. The magnetic beads were washed twice with ChIP buffer (20 mM Tris-HCl pH8.0, 150 mM NaCl, 2 mM EDTA, 1% Triton X-100), once with ChIP buffer including 500 mM NaCl, four times with RIPA buffer (10 mM Tris-HCl pH8.0, 0.25M LiCl, 1 mM EDTA, 0.5% NP-40, 0.5% Na·Deoxycholate), and once with TE buffer (pH 8.0). Chromatin was eluted twice from washed beads by adding elution buffer (20 mM Tris-HCl pH8.0, 100 mM NaCl, 20 mM EDTA, 1% SDS) and incubating for 15 min at 65°C. The crosslinking was reversed at 65°C for 6 hr and RNase A (Sigma) was added for 1 hr at 37°C followed by proteinase K (Ambion) treatment overnight at 50°C. ChIP-enriched DNA was purified using Phenol/Chloroform/Isoamyl alcohol extractions in phase-lock tubes. ChIP samples were end-repaired, A-tailed, and adaptor-ligated using QIAseq ultralow input library kit (Qiagen) according to the manufacturer’s instructions. Size selection using AMpure beads (Beckman Coulter) was performed to get 300-500 bp DNA and PCR amplification (8-10 cycles) were performed. Library quality and quantity were measured using TapeStation (Agilent) and Qubit (Thermo Fisher). We generated 50 bp paired-end reads on Illumina Hiseq 4000.

### Cell proliferation assay

We transduced cells with lentivirus carrying KRAB-dCas9 and sgRNA targeting candidate progrowth enhancers and performed puromycin (2 μg/ml) (InvivoGen) for three days postelectroporation to select against non-transduced cells. Seven days after the selection, the same number (4X10^5^ cells for HCT116; 2X10^5^ cells for A549) of viable cells was determined by trypan blue was split into 6-well plates triplicates. The number of viable cells was measured using trypan blue staining every 24 hours in an automated cell counter (Bio-rad) for three days.

### Proximity Ligation ChIP-sequencing (PLAC-seq)

PLAC-seq libraries were prepared for K562, HepG2, HCT116, and A549 cells as previously described with minor modifications (*35*). In brief, cells were cross-linked for 15 minutes at room temperature with 1% formaldehyde and quenched for 5 mins at room temperature with 0.2 M glycine (Thermo Fisher). The cross-linked cells were aliquot (~ 3X10^6^ cells) and resuspended in 300 μL lysis buffer (10mM Tric-HCl pH 8.0, 10mM NaCl, 0.2% IGEPAL CA-630) and incubated on ice for 15 minutes. The suspension was then centrifuged at 2,500 xg for 5 mins and the pellet was washed by resuspending in 300 μL lysis buffer and centrifuging at 2,500 xg for 5 mins. The pellet was resuspended in 50 μL 0.5% SDS and incubated for 10 mins at 62°C. 160 μL 1.56% Triton X-100 was added to the suspension and incubated for 15 mins at 37°C. 25 μl of 10X NEBuffer 2 and 100 U MboI were added to digest chromatin for 2 hours at 37°C with rotation (900 rpm). Enzymes were inactivated by heating for 20 mins at 62°C. Fragmented ends were biotin labeled by adding 50 μL of a mix containing 0.3 mM biotin-14-dATP, 0.3 mM dCTP, 0.3 mM dTTP, 0.3 mM dGTP, and 0.8 U μl^-1^ Klenow and incubated for 60 mins at 37°C with rotation (900 rpm). Ends were subsequently ligated by adding a 900 μL master mix containing 120 μL 10X T4 DNA ligase buffer (NEB), 100 μL 10% TritionX-100, 6 μL 20 mg ml^-1^ BSA, 10 μL 400 U μl^-1^ T4 DNA Ligase (NEB, high concentration formula) and 664 μL H2O and incubated for 120 mins at 23°C with 300 rpm slow rotation. Nuclei were pelleted for 5 mins at 4°C with centrifugation at 2,500 xg. For the ChIP, nuclei were resuspended in RIPA Buffer (10 mM Tris (pH 8.0), 140 mM NaCl, 1 mM EDTA, 1% Triton X-100, 0.1% SDS, 0.1% sodium deoxycholate) with proteinase inhibitors and incubated on ice for 10 mins. Sonication was performed using a Covaris M220 instrument (Power 75W, duty factor 10%, cycle per bust 200, time 10 mins, temperature 7°C) and nuclei were spun for 15 mins at 14,000 rpm at 4°C. 5% of supernatant was taken as input DNA. To the remaining cell lysate was added anti-H3K4me3 antibody-coated Dynabeads M-280 Sheep anti-Rabbit IgG (5 μg antibody per sample, Millipore, 04-745), followed by rotation at 4°C overnight for immunoprecipitation. The sample was placed on a magnetic stand for 1 min and the beads were washed three times with RIPA buffer, two times with high-salt RIPA buffer (10 mM Tris pH 8.0, 300 mM NaCl, 1 mM EDTA, 1% Triton X-100, 0.1% SDS, 0.1% deoxycholate), one time with LiCl buffer (10 mM Tris (pH 8.0), 250 mM LiCl, 1 mM EDTA, 0.5% IGEPAL CA-630, 0.1% sodium deoxycholate) and two times with TE buffer (10 mM Tris (pH 8.0), 0.1 mM EDTA). Washed beads were treated with 10 μg RNase A in extraction buffer (10 mM Tris (pH 8.0), 350 mM NaCl, 0.1 mM EDTA, 1% SDS) for 1 hour at 37°C, and subsequently 20 μg proteinase K was added at 65°C for 2 hours. ChIP DNA was purified with Zymo DNA clean & concentrator-5. For Biotin pull down, 25 μL of 10 mg ml^-1^ Dynabeads My One T1 Streptavidin beads was washed with 400 μl of 1X Tween Wash Buffer (5 mM Tris-HCl (pH 7.5), 0.5 mM EDTA, 1 M NaCl, 0.05% Tween) and supernatant removed after separation on a magnet. Beads were resuspended with 2X Binding Buffer (10 mM Tris-HCl pH 7.5, 1 mM EDTA, 2 M NaCl), added to the sample and incubated for 15 mins at room temperature. Beads were subsequently washed twice with 1X Tween Wash Buffer and in between heated on a thermomixer for 2 mins at 55°C with mixing and washed once with 1X NEB T4 DNA ligase buffer. Library prep was prepared using QIAseq Ultralow Input Library Kit (Qiagen). KAPA qPCR assay was performed to estimate concentration and cycle number for final PCR. Final PCR was directly amplified off the T1 Streptavidin beads according to the qPCR results, and DNA was size selected with 0.5X and 1X SPRI Cleanup and eluted in 1X Tris Buffer and paired-end sequenced.

### Analysis of tiling CRISPRi screen from the pilot study

The abundance of paired sgRNAs from D=0 and D=14 was mapped to the originally designed sequence using BWA (*66*). First, we analyzed the data using RELICS v1, a method specifically designed to analyze tiling CRISPR screens. RELICS v1 uses a Generalized Linear Mixed Model (GLMM) (*67*) to model gRNA counts across different pools. The output is a log Bayes Factor, which is calculated by comparing the ‘background’ model (a guide does not target a functional sequence) against a ‘functional sequence’ model (the guide targets a functional sequence). The functional model parameters are estimated by maximum likelihood from the observed guide counts for gRNAs targeting the MYC and PHB promoters (both are essential genes in cancer cells and used as positive controls in proliferation screen). In this study, the parameters of the background model were estimated by maximum likelihood from all the remaining guides. After computing the scores from both models, RELICS v1 calculates a RELICS score, also referred to as “CRISPRi score” in the proliferation-based screen, for each base pair by summing the log Bayes Factors of all paired gRNA overlapping genomic positions. A base pair is considered to be overlapped by gRNAs if it is within the ‘area of effect’ (AoE) for the CRISPR system used. We use 1kb as the AoE for a CRISPRi-based screen (*68*). We selected genomic regions with an averaged CRISPRi score above 5 ad candidate pro-growth enhancers. This threshold was chosen to correspond to FDR < 0.1 based on simulated data from CRSsim (*34*). To generate high confidence candidate pro-growth enhancers across different cell types, we further selected the reproducible candidate enhancers, which were identified from two independent pipelines, RELICS v1 and CRISPY. CRISPY is an improved pipeline developed from our previous study (*62*) and allows the flexibility of processing different types of screening data, data quality control, and peak calling for positive elements. RELICS v1 can be obtained from GitHub (https://github.com/patfiaux/RELICS/releases/tag/v1.0), and we used default settings for the analysis in this study. CRISPY can be obtained from GitHub (https://github.com/MichaelMW/crispy), and we performed the analysis with parameter -n 0.05, which only outputs the candidate peaks with FDR < 0.05.

### Analysis of genome-wide CRISPRi screen

The abundance of paired sgRNA from D=0 and D=14 was mapped to the originally designed sgRNA sequence using BWA (*66*). We used MAGeCK (*43*) to perform data quality assessment and identify candidate pro-growth enhancers. For data quality assessment, we used the MAGeCK-mle module to calculate β-score (guide RNA depletion index) between biological replicates from the screen in two individual clones. A negative β-score indicates a target is negatively selected. Pearson correlation was performed to determine the correlation between β-score from two individual clones. Next, we used the RRA algorithm to obtain the RRA score for the individual target gene or H3K27ac peak. Negative control sgRNAs were provided to generate the null distribution when calculating the *P* values. The threshold value for candidate pro-growth enhancers is FDR < 0.2 based on the ROC curve with 80% sensitivity. The information of selected candidate pro-growth enhancers is listed in Table S7.

Removal of candidate pro-growth enhancers targeting by low-specificity sgRNA pairs To evaluate the potential of off-target sgRNA-mediated toxicity to affect cellular proliferation, we computed a specificity score for all sgRNAs included in the genome-wide screen using GuideScan (*69*) from the webtool. For every paired gRNA, we used the lowest specificity score from one of the two sgRNAs for the representation. A previous study (*70*) has shown that sgRNAs with specificity score < 0.2 tend to have substantial off-target toxicity in CRISPR based screens. To minimize false-positive hints by off-target toxicity in our screen, we removed candidate pro-growth enhancers targeted by more than two sgRNA pairs (40%) with specificity score < 0.2 or with unidentified specificity score. Total we further removed 70 candidate progrowth enhancers targeted by low-specificity sgRNA pairs (Table S8).

### ChIP-seq data processing

Each fastq file was aligned to the human genome (hg38) with bowtie2 (Version 2.3.4.3) (*77*). SAMtools (Version 1.9) (*72*) and MarkDuplicates (Picard) were used to filter (MAPQ < 30) and clean data post alignment. The reads were then converted to reads per kilobase per million in 200 bp bins using deepTools2 (Version 3.5.0) (*73*). We obtained reproducible H3K27ac peaks in HCT116 from ENCODE and filtered out H3K27ac peaks located within 3-kb around annotated TSS using bedtools. For each distal H3K27ac peaks (n=22,744), we obtained ChIP-seq fold enrichment over input in each sample using bigWigAverageOverBed (https://github.com/ENCODE-DCC/kentUtils/blob/master/bin/linux.x86_64/bigWigAverageOverBed). Pearson correlation was performed to determine the correlation of H3K27ac signal between different samples.

### Analysis of key epigenetic features with pro-growth enhancer

For key epigenetic features analysis, each functional enhancer peak was then scored using readdepth normalized signals from DNase-seq, fold-change over input from ChIP-seq data, and the fitness score from tiling screen using bigWigAverageOverBed to extract averaged signal for progrowth enhancer peaks across different cell types. We used spearman’s rank correlation to determine the correlation between key epigenetic features and pro-growth enhancers across multiple cell types.

### PLAC-seq data processing

PLAC-seq data was processed with MAPS (*44*) to normalize reads and identify long-range chromatin interactions. Specifically, MAPS aligned raw paired-end reads with BWA (*66*) to the reference genome hg38. Uniquely mapped reads were kept and split into intra-chromosomal reads and inter-chromosomal reads. We used the following steps to identify intra-chromosomal chromatin interactions. Each chromosome was first divided into 10 kb bins. Histone H3K4me3 ChIP-seq peaks (downloaded from ENCODE) were used as the anchor, and 10 kb bins overlapping with these ChIP-seq peaks were defined as the anchor bin. Depending on whether none, one, and two bins are the anchor bin, 10 kb bin pairs were further defined as ‘NOT’, ‘XOR’, and ‘AND’ sets. Only bin pairs in the ‘XOR’ and ‘AND’ sets were kept for downstream analysis. The raw contact frequency between two 10 kb bins in the ‘XOR’ and ‘AND’ sets was then fitted into a zero-truncated Poisson model to obtain the normalized contact frequency. Significant interactions were identified with FDR corrected p-value cutoff of 0.01. Significant interactions were further grouped into clusters if two interactions were within 10 kb.

### Differential chromatin interaction analysis

For differential interaction analysis in H3K4me3 PLAC-seq, the raw contact counts in 10 kb resolution bins were used as inputs, and we stratified the inputs into every 10-kb genomic distance to minimize the bias from genomic distance. Since each input showed negative binomial distribution, we used edgeR (*74*) to get the initial set of differential interactions. Next, we removed bins with less than 20 contact counts in each sample of two replicates from the downstream analysis. To avoid the antibody bias and TSS with differential H3K4me3 level, we further removed chromatin contacts overlapping with differential H3K4me3 ChIP-seq peaks at TSS (with fold-change decreased more than 50% than in HCT116). We only compared the chromatin contacts between different cell types at TSS of the predicted target genes with the same level of H3K4me3 ChIP-seq peaks.

### Target gene prediction of pro-growth enhancer

We used functional similarity and chromatin interaction to identify the targets of pro-growth enhancers. From the tiling screen, we demonstrated cellular proliferation assay usage to identify strong functional enhancer for *MYC* oncogene across multiple cell types and found that the distance between a majority of functional enhancer and target gene is within 500-kbp. Thus, we first obtained a gene dependence score from CRISPR-Cas9 knockout screen in HCT116 (*26, 39*). We first utilized chromatin interactions generated from H3K4me3 PLAC-seq to identify 155 target genes that spatially interact with 115 pro-growth enhancers and are also important for fitness. In addition, we also extended 500-kbp upstream and downstream of pro-growth enhancers and identified all essential genes located within that range. Overall, we identified 755 E-G pairs from 278 pro-growth enhancers using this approach. Together, we predicted 910 E-G pairs from 393 pro-growth enhancers (80.5% of total pro-growth enhancers identified in the screen). The predicted E-G pairs are listed in Table S9.

### Survival analysis

Clinical outcome data and normalized ATAC-seq counts were obtained from UCSC Xena (*75*). Cox proportional hazards regression model and the hazard ratio was computed in R using package survival. Package survminer was used for drawing the Kaplan-Meier plots and defining the optimal threshold (surv_cutpoint). For gene expression (*MYC, COASY, WDR18*) survival analysis in 398 colon cancer samples, clinical outcome data, and corresponding normalized RNA-seq data were obtained from UCSC Xena (*75*). Package survminer was used for drawing the Kaplan-Meier plots and defining the optimal threshold (surv_cutpoint) using the maximally selected rank statistics (*76, 77*).

### Motif enrichment analysis

To identify potential regulators of pro-growth enhancers, we performed motif analysis using the AME utility (5.1.15.1.1) of the MEME suite (*78*). For enrichment of known motifs, one-tailed Fisher’s exact test was used to calculate significance. We used 11,111 distal H3K27ac peaks as the background and the default setting, *E*-value (the motif *P*-value multiplied by the number motifs in the input) cutoff of < 10, was chosen for known motifs from HOCOMOCO database (HOCOMOCO v11 Full) (*79*). The motifs with significant *P* values (-log_10_ adj p-value >2) were selected and reported.

### Analysis of key transcription factor

We downloaded 790 DNase-seq datasets (Table S10) from the ENCODE portal. We then applied the Taiji pipeline (*49*) to rank transcription factors. For each sample we constructed the TF regulatory network by scanning TF motifs at the accessible chromatin regions and linking them to the nearest genes. The network is directed with edges from TFs to target genes. The genes’ weights in the network were determined based on the relative accessibility of their promoters. The weights of the edges were calculated by the relative accessibility of the promoters of the source TFs. We then used the personalized PageRank algorithm to compute the ranking scores for the TFs in the network.

### Data source

We downloaded DNase-seq and ChIP-seq data generated by the ENCODE Project Consortium and Roadmap Epigenomics Project. Gene expression and gene dependence data were obtained from the Cancer Cell Line Encyclopedia and the Cancer Dependency Map Project. All data used in each figure are listed in Table S11.

### Software for data analysis and graphical plots

We used the following software for data analysis and graphical plots: R Bioconductor (version 3.6.1), edgeR (version 3.12), Survival (version 3.2-7), Survminer (version 0.4.8), BEDTools (*80*), Integrative Genomics Viewer (version 2.4.10) (*81*), MAGeCK (version 0.5.9.2) (*43*), RELICS v1 (https://github.com/patfiaux/RELICS/releases/tag/v1.0), CRISPRY (https://github.com/MichaelMW/crispy), MAPS (https://github.com/ijuric/MAPS) (*44*), bigWigAverageOverBed (https://github.com/ENCODE-DCC/kentUtils/blob/master/bin/linux.x86_64/bigWigAverageOverBed).

### Genome build

All coordinates are reported in human genome build hg38.

---

**Fig. S1.**
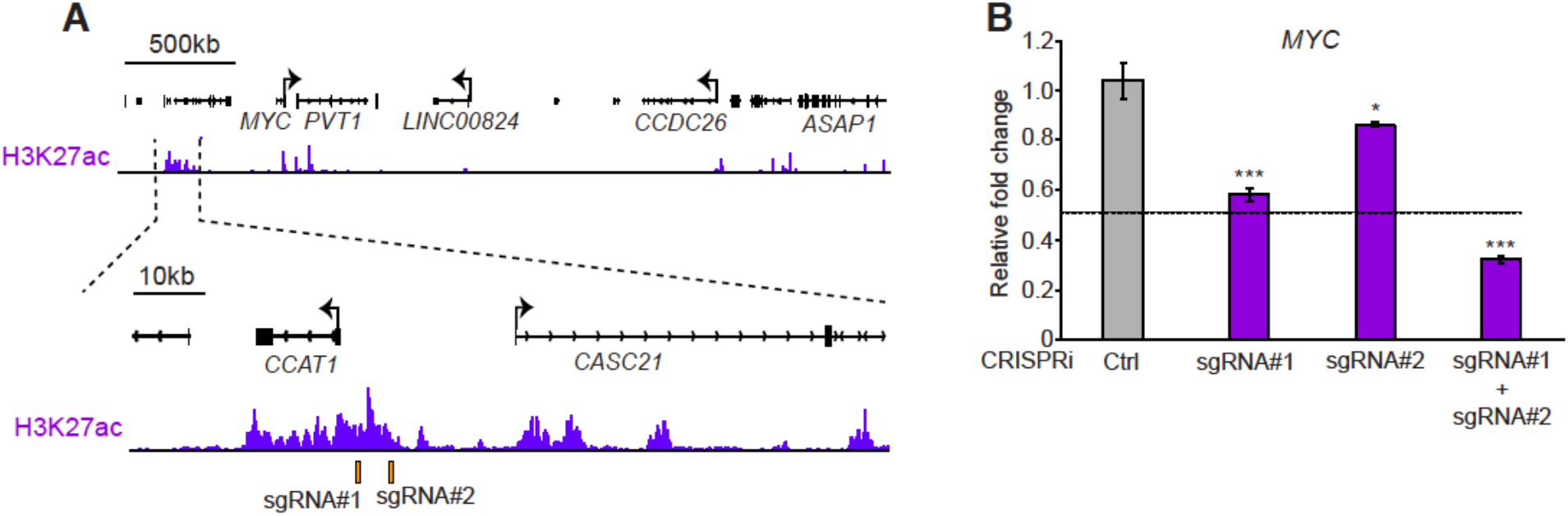
Dual sgRNAs design improves the efficiency of epigenetic silencing mediated by KRAB-dCas9 at super-enhancer. (**A**) Selection of sgRNAs targeting a previous reported superenhancer upstream of *MYC* locus in HCT116. The orange bar presents the target site of each sgRNA with tracks showing H3K27ac signal at this selected genomic locus. (**B**) Greater gene repression achieved using dual sgRNAs targeting super-enhancer. Messenger RNA expression level was measured using RT-qPCR after targeting different genomic loci with KRAB-dCas9. Data shown are mean ± SD of three technical replicates from one representative experiment of two biological replicates performed. *P*-values were determined by two-tailed Student’s *t*-test (* and *** indicates *p* < 0.1 and *p* < 0.001 respectively). The dashed line indicates the expected fold-change by targeting two genomic loci with KRAB-dCas9.

**Fig. S2.**
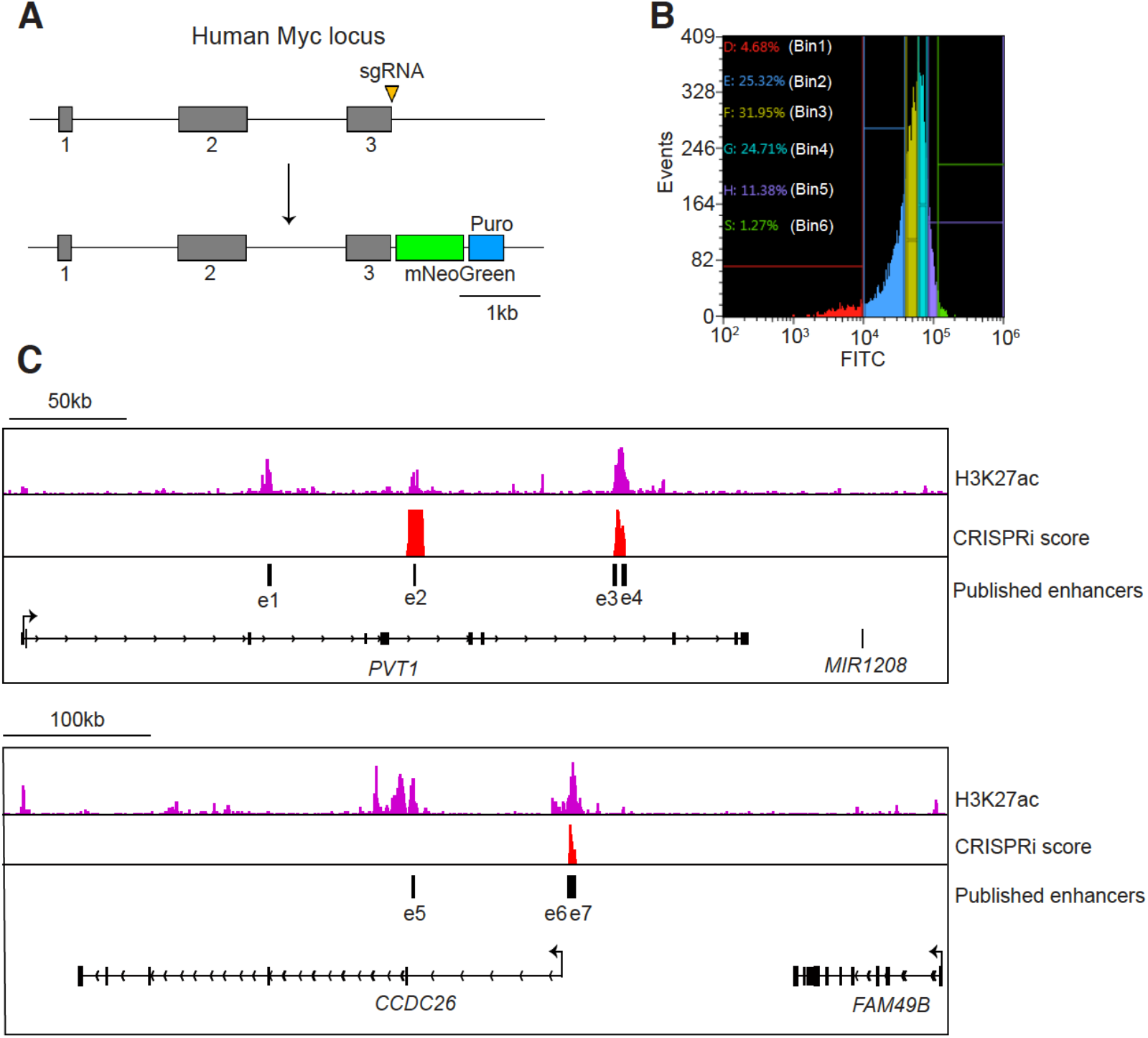
Examination of the performance of tiling paired sgRNA library using CRISPRi to identify functional enhancers for *MYC* oncogene in K562 cells. (A) Scheme of the experimental strategy to illustrate CRISPR/Cas-9 mediated gene targeting for knocking in a fluorescent reporter, mNeoGreen, into C-terminal of the *MYC* gene in K562 cells. The yellow triangle represents the sgRNA target site at the last exon of *MYC*. (B) K562 cells were divided into 6 bins and sorted using FACS based on the expression level of mNeoGreen (FITC) after lentivirus transduction carrying KRAB-dCas9 tiling paired sgRNA library. (C) The functional enhancers identified from tiling paired sgRNA screen. A close-up view of the known functional enhancers identified from the previous study (*20*), with tracks showing H3K27ac ChIP-seq signal and CRISPRi scores generated from tiling paired sgRNAs screen in selected genomic loci.

**Fig. S3.**
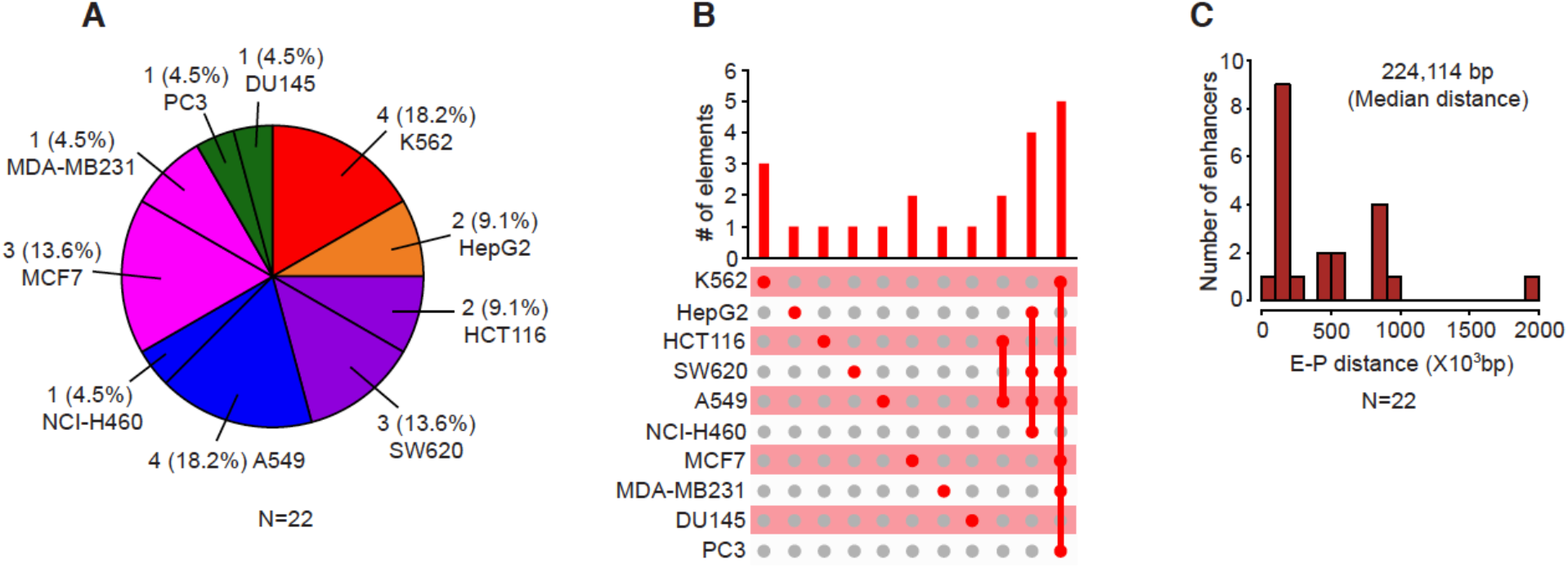
Summary of pro-growth enhancers identified from the proliferation-based CRISPRi pooled screen. (A) The number of pro-growth enhancers identified from each cell line. (B) The number of cell-type-specific and pan-cancer pro-growth enhancers. Red dots and gray dots indicate the presence and absence of pro-growth enhancers, respectively. (C) The distance between pro-growth enhancers and the TSS of target genes.

**Fig. S4.**
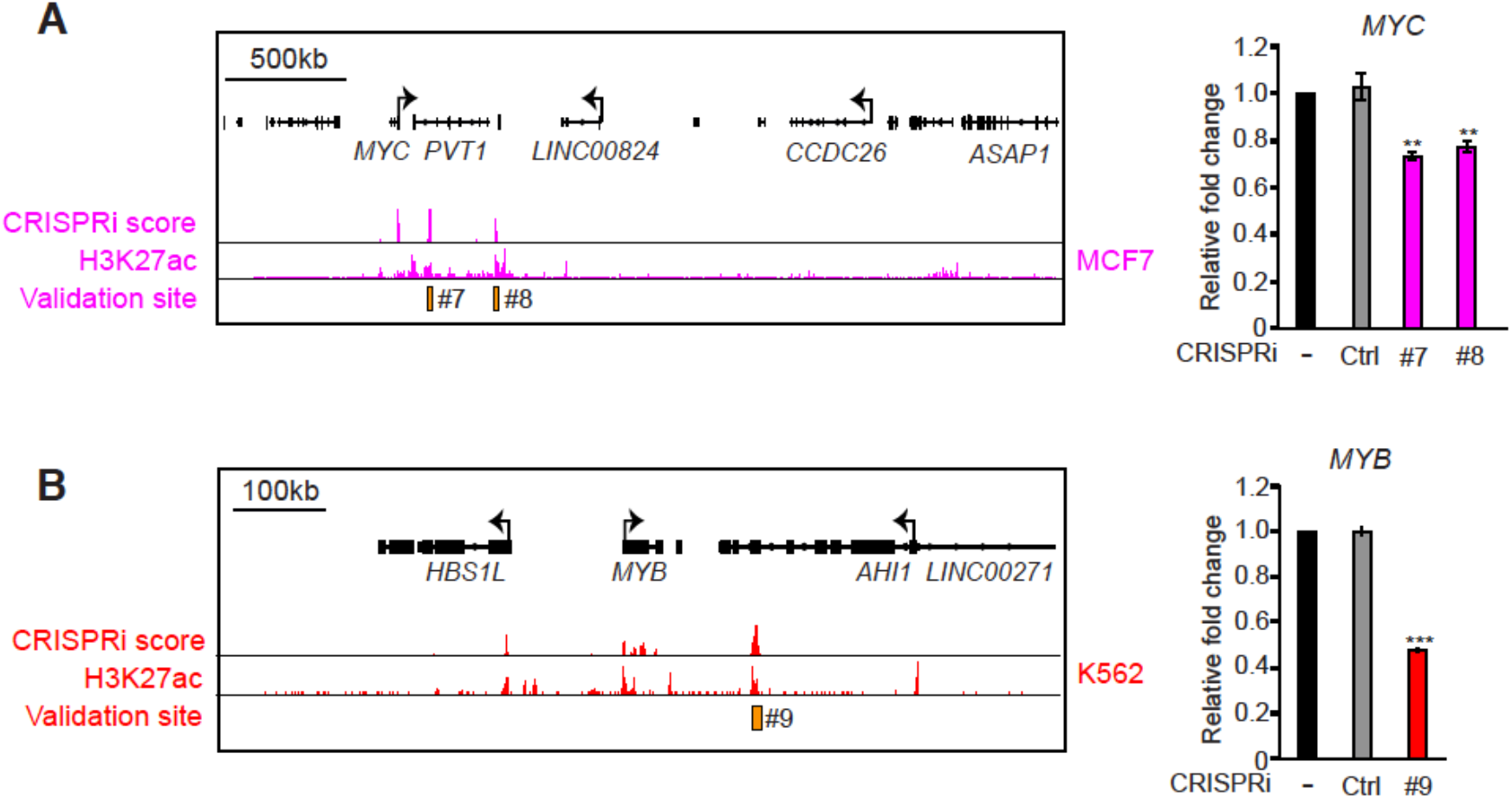
Pro-growth enhancers are important for the expression of key oncogenes. (A-B) Left: Pro-growth enhancers identified in MCF7 (A) and K562 (B). Right: The measurement of relative fold change in *MYC* (A) and *MYB* (B) oncogenes by RT-qPCR after epigenetic silencing pro-growth enhancers using KRAB-dCas9. Data shown are mean ± SD of three technical replicates from one representative experiment of two biological replicates performed. *P*-values were determined by a two-tailed Student’s *t*-test (** and *** indicates *p* < 0.01 and *p* < 0.001 respectively).

**Fig. S5.**
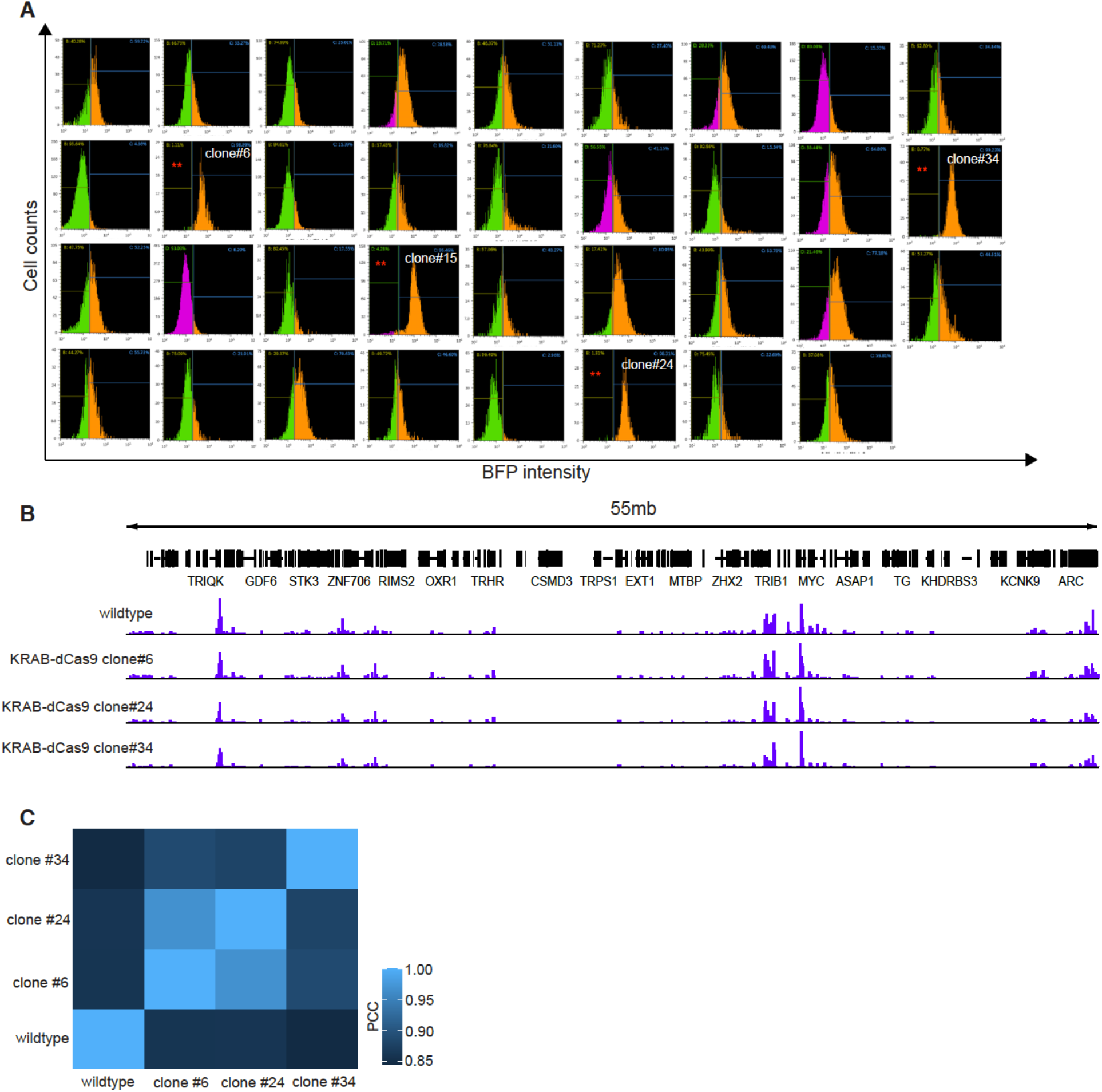
H3K27ac signal in KRAB-dCas9 stably expressed HCT116 cell clones is similar to parental cells. (A) The selection of KRAB-dCas9 clones based on the expression level of BFP (dCas9-TagBFP-KRAB) by FACS analysis. (B) Genome browser snapshot comparing H3K27ac ChIP-seq signal between parental HCT116 (wildtype) and three KRAB-dCas9 stable clones (KRAB-dCas9 clone #6, #24, #34) within the selected 55-Mbp genomic locus. (C) Correlation of H3K27ac signal in distal enhancers between parental HCT116 (wildtype) and three KRAB-dCas9 stable clones (KRAB-dCas9 clone #6, #24, #34).

**Fig. S6.**
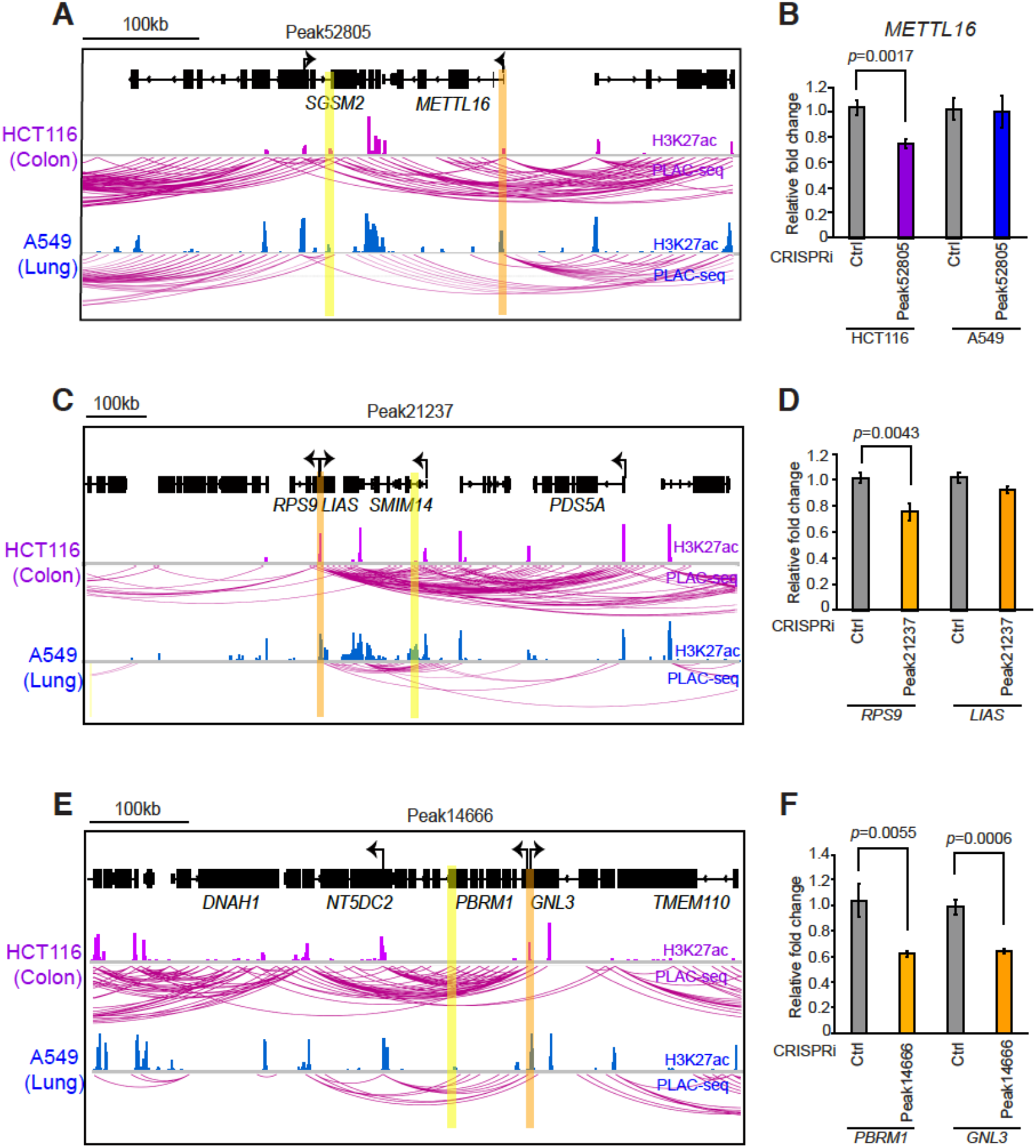
Validation of multiple predicted E-G pairs. (A, C, E) Comparison of chromatin interactions identified by H3K4me3 PLAC-seq in the indicated genomic regions from HCT116 and A549. The pro-growth enhancers are highlighted in yellow and the predicted target genes are highlighted in orange. (B, D, F) Relative RNA levels of predicted target gene determined by RT-qPCR after targeting candidate pro-growth enhancer with KRAB-dCas9 (n=3 per group). Data shown are mean ± SD of three technical replicates from one representative experiment of two biological replicates performed. *P*-values were determined by a two-tailed Student’s *t*-test.

**Fig. S7.**
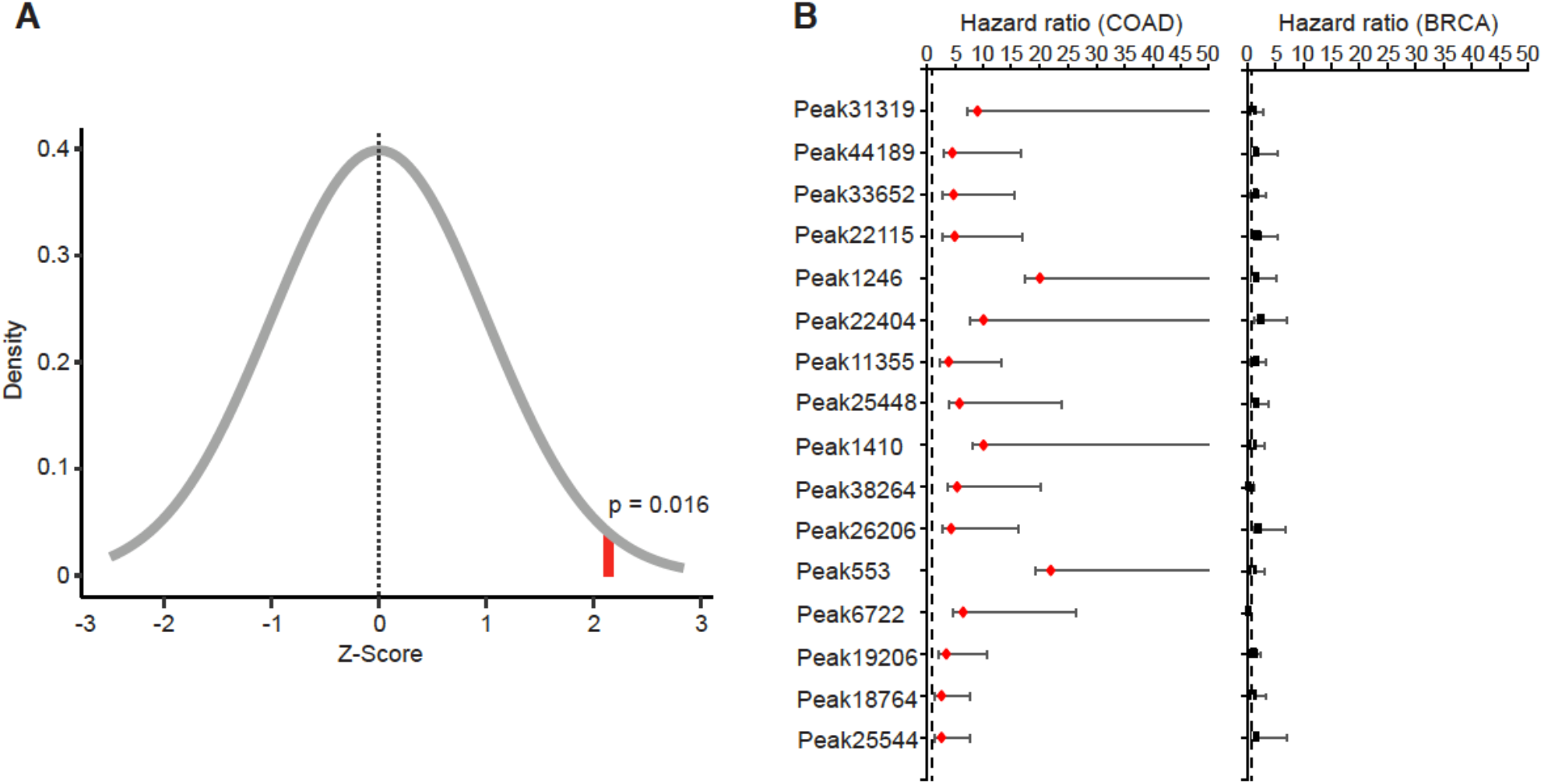
Chromatin accessibility of pro-growth enhancers is not associated with poor clinical outcome in breast cancer samples. (A) Enrichment of pro-growth enhancers associated with poor clinical outcome in COAD samples. The distribution shown is the result of 528 randomly selected ATAC peaks that overlapped with distal putative enhancers in HCT116 after repeating a total of 35 randomized selections. The red bar indicates the observed number of progrowth associated with higher hazard ratio. *P*-values was determined by the one-tailed one-sample t-test. (B) Cox proportional hazards regression analysis of ATAC-seq peaks overlapping with pro-growth enhancers in 38 colon cancer samples (left) and 63 breast cancer samples (right). Error bars represent 95% confidence intervals. Dashed lines indicates where the hazard ratio equals one. COAD, colon adenocarcinoma; BRCA, breast invasive carcinoma.

**Fig. S8.**
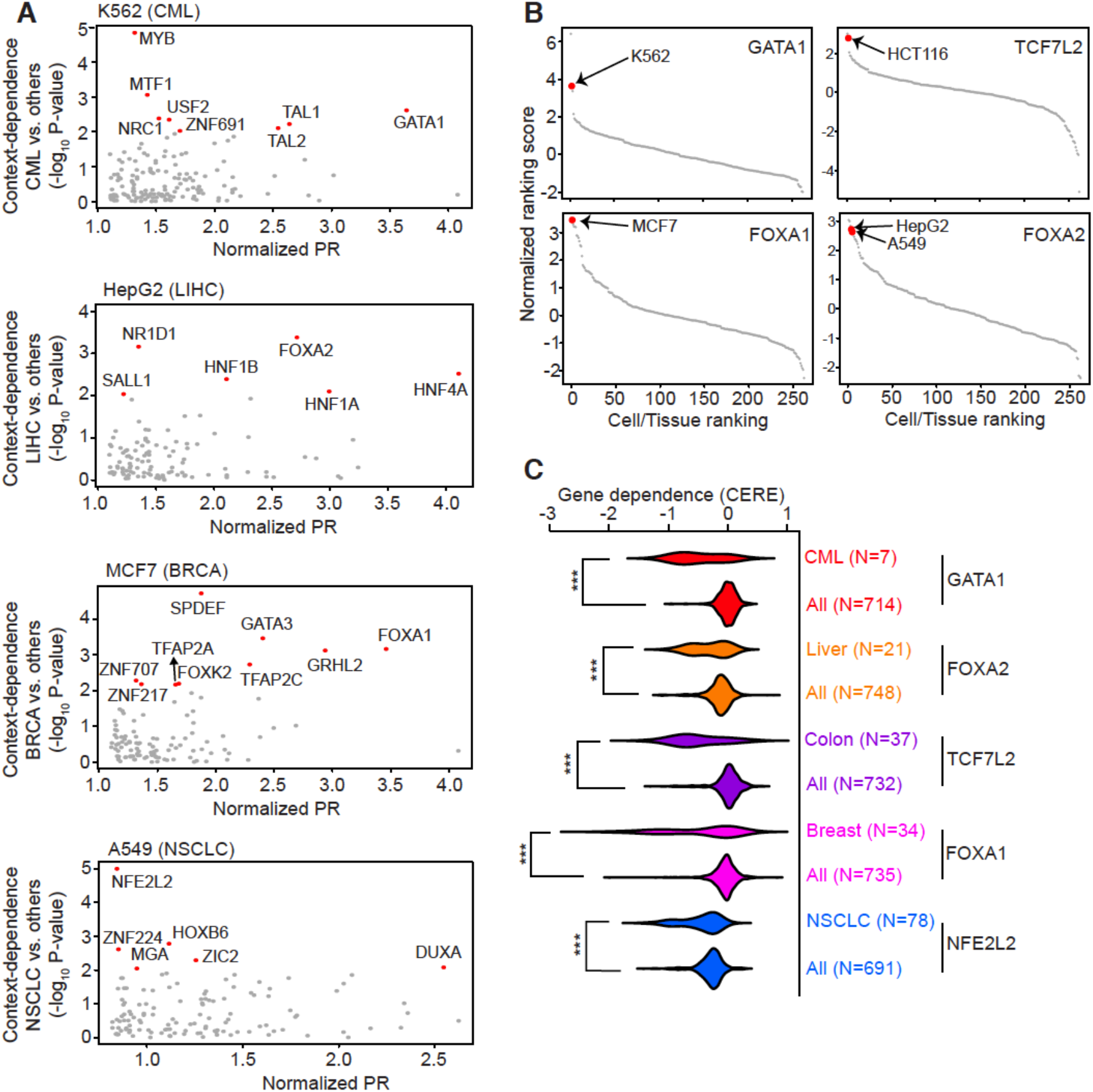
Identification of critical transcription factor to regulate pro-growth enhancers. (A) Identification of candidate TFs from four different cancer types using normalized page-rank score (normalized PR) and gene dependency. The red dots represent selected candidate TFs with high normalized page-rank score and significant context-dependence (p < 0.01). (B) The distribution of ranking score from lineage-specific TFs in 262 human cell lines and tissues. Red dots indicate the cell lines that are included in this study. (C) Gene dependence (CERE) of lineage-specific TFs in human cancer cell lines (***: p < 0.001). *P*-values were determined by two-side Wilcoxon test. CML, chronic myelogenous leukemia; LIHC, liver hepatocellular carcinoma; BRCA, breast invasive carcinoma; NSCLC, non-small-cell lung carcinoma.

**Fig. S9.**
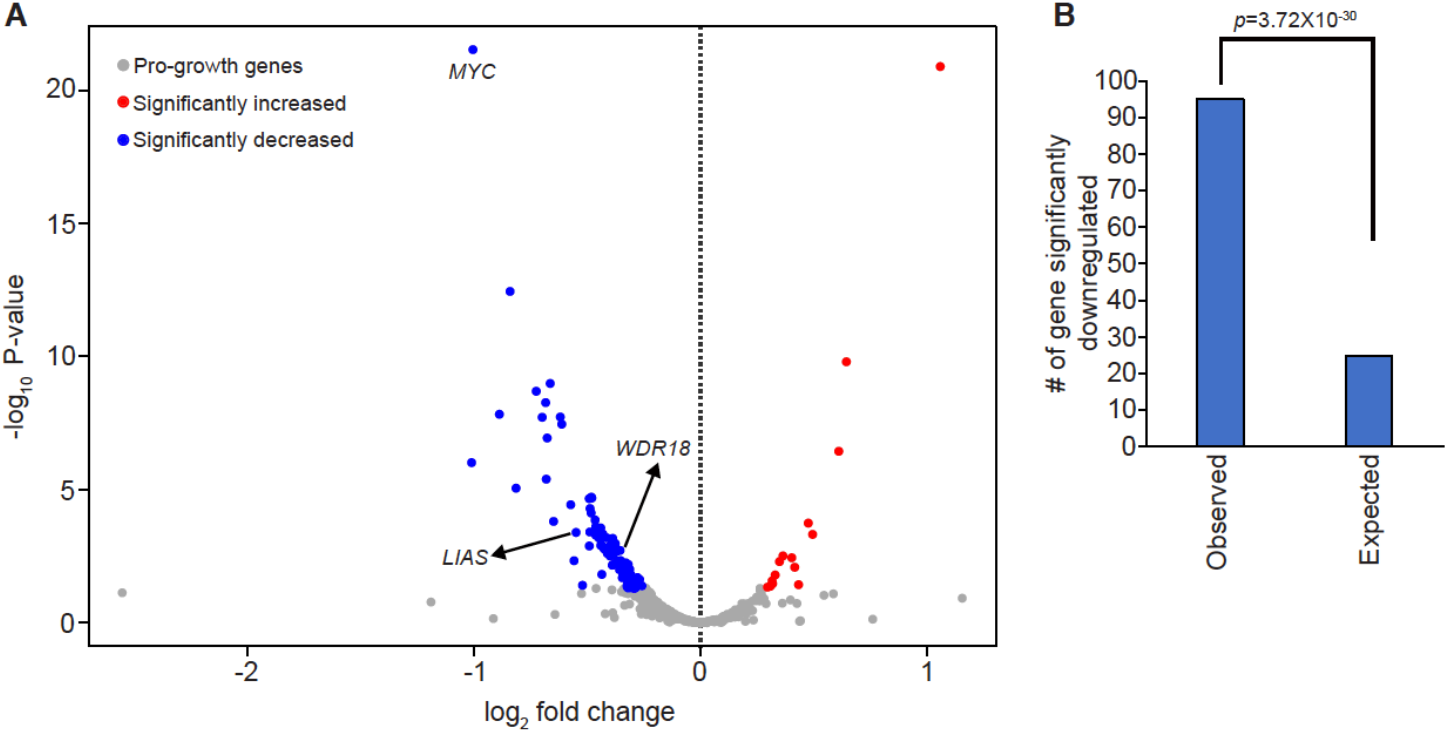
RNA-seq analysis of pro-growth gene expression in *Tcf7l2* knockout cells. (A) Volcano plot showing the changes in gene expression of pro-growth gene after the loss of *Tcf7l2* in HCT116 cells. The red dots represent significantly upregulated genes, the blue dots represent significantly downregulated genes (adjusted p-value <0.05), and the gray dots represent insignificant differentially expressed genes. (B) Comparison between the number of significantly downregulated pro-growth genes and randomly selected pro-growth enhancers. *P*-value was determined by a hypergeometric test.

**Table S1 (separate file).**

**Tiling CRISPRi paired sgRNA library sequences and screening data.**

Sequences, annotations, and raw counts for paired sgRNA library.

**Table S2 (separate file).**

**Genome-scale CRISPRi paired sgRNA library sequences and screening data.**

Sequence, annotations, and raw counts for genome-scale sgRNA library.

**Table S3 (separate file).**

**Oligo information for sequencing library preparation from CRISPRi screen.**

Primer sequence and index information for 1^st^ and 2^nd^ round PCR.

**Table S4 (separate file).**

**Sequences of guide RNA for single guide RNA construct.**

**Table S5 (separate file).**

**Sequences of guide RNA for dual guide RNA construct.**

**Table S6 (separate file).**

**Primer sequences for RT-qPCR.**

**Table S7 (separate file).**

**Candidate pro-growth enhancers identified from genome-scale CRISPRi screen in HCT116 cells.**

Annotations and statistical information (p-value, FDR, and log_2_ fold-change) of pro-growth enhancer.

**Table S8 (separate file).**

**Summary of sgRNA pairs with low specificity score targeting to pro-growth enhancers.**

**Table S9 (separate file).**

**Summary of predicted enhancer and target gene pairs.**

Annotation of pro-growth enhancers and their target genes based on chromatin interaction information (500kb around TSS of target gene or chromatin contacts identified by PLAC-seq).

**Table S10 (separate file).**

**Information of DNase-seq datasets analyzed in Taiji pipeline.**

Cell/ Tissue types and accessory number of DNase-seq datasets.

**Table S11 (separate file).**

**Source for epigenomic data utilized in each figure.**

## Notes

### Competing Interest Statement

B.R. is a co-founder and consultant for Arima Genomics, Inc., and a co-founder of Epigenome Technologies, Inc.

